# *Insm1*-expressing neurons and secretory cells develop from a common pool of progenitors in the sea anemone *Nematostella vectensis*

**DOI:** 10.1101/2021.04.09.439178

**Authors:** Océane Tournière, Henriette Busengdal, James M. Gahan, Fabian Rentzsch

## Abstract

Neurons are highly specialized cells present in nearly all animals, but their evolutionary origin and relationship to other cell types is not well understood. We use here the sea anemone *Nematostella vectensis* as a model system for early-branching animals to gain fresh insights into the evolutionary history of neurons. We generated a transgenic reporter line to show that the transcription factor *NvInsm1* is expressed in post-mitotic cells that give rise to various types of neurons and secretory cells. Expression analyses, double transgenics and gene knockdown experiments show that the *NvInsm1*-expressing neurons and secretory cells derive from a common pool of *NvSoxB(2)*-positive progenitor cells. These findings, together with the requirement for *Insm1* for the development of neurons and endocrine cells in vertebrates, support a close evolutionary relationship of neurons and secretory cells.

## INTRODUCTION

Neurons are cells with a high diversity of morphological, molecular and physiological properties. The common theme of neurons is the presence of subcellular structures that allow fast and precise intercellular communication. Many neurons have long processes, collectively termed neurites, that are used for the propagation of electrical excitation. These electrical signals trigger the release of the content of small vesicles either in a highly specific manner at chemical synapses or with lower temporal and spatial resolution via neurosecretion. The sophisticated molecular machineries that enable electrical conductance and vesicle release resemble those found in other cell types. The electrical excitability and conduction in muscle cells is based on the same types of voltage-gated sodium and calcium channels as that of neurons, whereas the release of synaptic vesicles involves many proteins associated with the vesicle and plasma membrane, respectively, that also function in vesicle release from non-neural secretory and endocrine cells. The evolution of neurons thus likely resulted from the elaboration of different pre-existing cellular modules (Arendt, 2020; Brunet and Arendt, 2016; Burkhardt and Sprecher, 2017; Kristan, 2016; Liebeskind et al., 2017; Moroz, 2009; Varoqueaux and Fasshauer, 2017). Since the presence of the constituents of these different cellular modules within the same cell requires their co-ordinated expression, analysing the gene regulatory program and the developmental origin of neurons in different species can provide valuable insights into the evolutionary relationship between neurons and other cell types. Here we follow this approach by addressing the development of neurons and gland/secretory cells in the sea anemone *Nematostella vectensis*, a member of an animal clade that separated from the bilaterian lineage (e.g. chordates, arthropods, nematodes, annelids) more than 600 million years ago (dos Reis et al., 2015; Park et al., 2012).

Vertebrate neurons originate from the ectoderm whereas endocrine cells of the gastrointestinal system originate from the endoderm. Despite this significant difference, there are many similarities in the development of neurons and endocrine cells. They involve the use of conserved signalling molecules and transcription factors like Notch signalling, soxB genes and bHLH genes of the atonal, neurogenin and achaete scute families (Arntfield and van der Kooy, 2011; Ernsberger, 2015; Hartenstein et al., 2017). For example, inactivation of Notch signalling in mice results in a neurogenic phenotype characterized by an excess of neurons (de la Pompa et al., 1997; Lutolf et al., 2002) and in the development of excess endocrine cells in the pancreas (Apelqvist et al., 1999). Similarly, transcription factors of the neurogenin family are required for the development of olfactory sensory neurons (Cau et al., 2002) and for the formation of endocrine cells in the gastrointestinal tract (Gradwohl et al., 2000; Jenny et al., 2002; Lee et al., 2002). Interpretations of these similarities are confounded, however, by the use of different paralogs in different cell types and tissues, e.g., *neurogenin1* in the olfactory epithelium and *neurogenin3* in the gastrointestinal tract, and by the widespread functions of some of these gene families in the specification of other cell types (e.g. (Buas and Kadesch, 2010; Pajcini et al., 2011).

A more striking case is the zinc finger transcription factor Insulinoma-associated protein 1 (Insm1 or IA-1). During mouse embryonic development, *Insm1* is expressed broadly in the central and peripheral nervous system, in the stomach, intestine, pancreas, thymus, thyroid and adrenal glands (Breslin et al., 2003; Duggan et al., 2008; Gierl et al., 2006; Mellitzer et al., 2006). Analyses of knock-out mice revealed that *Insm1* is required for the development of neurons throughout the brain (Farkas et al., 2008; Jacob et al., 2009) and of neural crest-derived cells of the sympatho-adrenal lineage in the peripheral nervous system (Wildner et al., 2008). Moreover, *Insm1* controls the development of endocrine cells in the pancreas, lung, intestine and pituitary gland (Gierl et al., 2006; Jia et al., 2015b; Mellitzer et al., 2006; Welcker et al., 2013). The broad roles in the development of neurons and endocrine cells make *Insm1* a particularly interesting candidate gene for addressing the evolutionary relationship between neural and secretory cells.

Cnidarians (e.g. sea anemones, corals, jellyfish) are the sister taxon to bilaterians (Dunn et al., 2014; Telford et al., 2015) and the earliest-branching group of animals whose nervous system unambiguously shares a common origin with that of bilaterians (Hartenstein and Stollewerk, 2015). There are two major groups of cnidarians, anthozoans and medusozoans, with the polyp as the body form common to both groups and the medusa as a pelagic life cycle stage present only in medusozoans. Polyps can be described as a tube with one opening, that is surrounded by tentacles used for the capture of prey (Technau and Steele, 2011). Notwithstanding local differences in the density of neurons and neurites, the nervous system of cnidarian polyps is best described as a nerve net that lacks brain-like centralization. Three general classes of neural cells are distinguished in cnidarians: sensory/sensory-motor neurons, ganglion cells (potentially equivalent to interneurons) and cnidocytes, the cnidarian-specific stinging cells (Galliot and Quiquand, 2011; Rentzsch et al., 2017; Watanabe et al., 2009a). The developmental origin of neural cells appears to differ between different groups of cnidarians. In hydrozoans (a subgroup of medusozoans), so-called interstitial stem cells give rise to neural cells. Interestingly, the interstitial stem cell population also gives rise to gland/secretory cells and gametes, but similar non-epithelial stem cells have not been observed outside this particular group of cnidarians (Bosch et al., 2010; Frank et al., 2009; Watanabe et al., 2009a; Watanabe et al., 2009b). The molecular control of neurogenesis in cnidarians is currently best understood in the anthozoan *Nematostella vectensis* (Layden et al., 2016; Rentzsch et al., 2017), in which soxB genes, Notch signalling and bHLH genes of the atonal/neurogenin and achaete scute families regulate the development of neural cells (Layden et al., 2012; Layden and Martindale, 2014; Richards and Rentzsch, 2014; Richards and Rentzsch, 2015; Watanabe et al., 2014). In particular, a pool of epithelial *NvSoxB(2)*-expressing progenitor cells has been identified that gives rise to sensory neurons, ganglion cells and cnidocytes during embryonic development (Richards and Rentzsch, 2014). Inhibition of *NvSoxB(2)* or *NvAtonal-like* (*NvAth-like*) resulted in a strong reduction in the number of neurons and cnidocytes, whereas inhibition of Notch signalling increased their numbers (Layden and Martindale, 2014; Richards and Rentzsch, 2014; Richards and Rentzsch, 2015). These observations identified a substantial degree of similarity in the genetic program regulating the early stages of *Nematostella* and vertebrate neurogenesis. In contrast to vertebrates, however, neurogenesis in *Nematostella* occurs both in the ectoderm and in the mesendoderm, the only two germ layers in the diploblastic cnidarians (Nakanishi et al., 2012; Richards and Rentzsch, 2014). The development of gland/secretory cells is not well understood in *Nematostella*, but it has recently been shown that gland/secretory cells expressing either digestive enzymes or insulin-like peptides are located in tissues of ectodermal origin. These tissues are the ectodermal part of the pharynx and the septal filaments, which are extensions of the pharyngeal ectoderm (Steinmetz et al., 2017). The septal filaments extend along the oral-aboral axis of the animals and are part of internal structures of anthozoans called mesenteries **(Figure 1A)**. The ectodermal origin of these gland/secretory cells is in contrast to the endodermal origin of equivalent cells in the vertebrate midgut and pancreas (Steinmetz et al., 2017).

**Figure 1:**
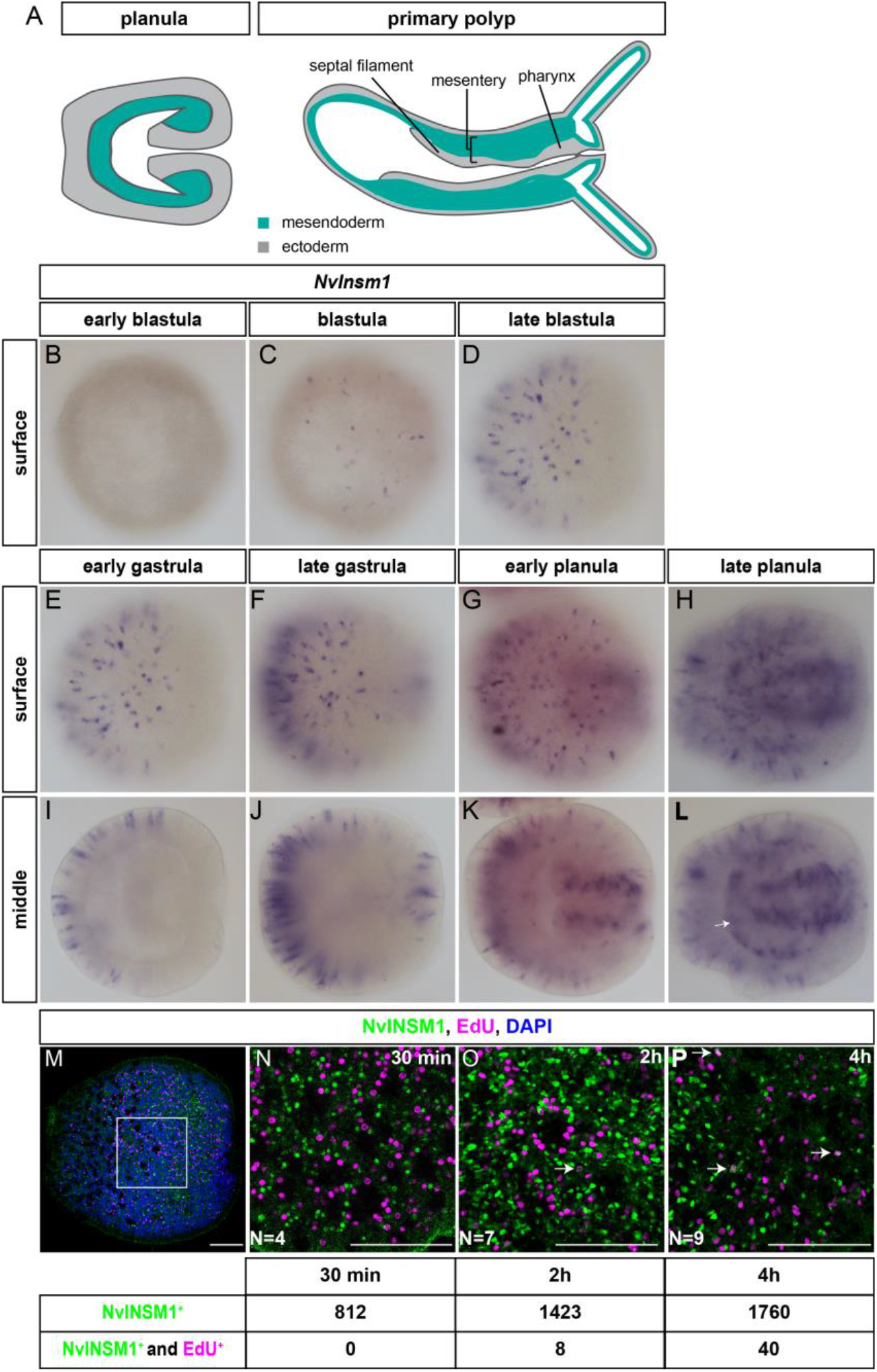
*NvInsm1* is expressed in scattered post mitotic cells. (A) Cartoons of *Nematostella* at planula and primary polyp stage. Tissues of mesendodermal and ectodermal origin are labaled in green and grey, respectively. Aboral pole to the left, drawings are not to scale. (B-L) *NvInsm1 in situ* hybridization with the developmental stage indicated on the top. Mid-lateral views with the aboral pole to the left. (B-H) are focused on the surface of the specimens, (I-L) are optical sections at the middle of the specimens. Starting at blastula stage (12hpf), *NvInsm1* is expressed in scattered single cells mainly in the aboral half of the embryos. At early planula stage *NvInsm1* starts being expressed in the forming pharynx (K). At late planula stage (L), *NvInsm1* starts being expressed in the mesendoderm, indicated by the white arrow. (M-P) Immunostaining of NvINSM1 (green), EdU (magenta) and DAPI (blue) at planula stages. Staining is indicated at the top. (N) None of the ectodermal cells expressing NvINSM1 incorporated EdU after a 30min pulse. After 2H (O) and 4H (P) of EdU exposure, some EdU^+^/NvINSM1^+^ cells can be detected (white arrows in O, P). N stands for the number of embryos analyzed. Scale bar represents 20 μm.

To better understand the relationship between neurons and gland/secretory cells, we studied here the role of *NvInsm1* in *Nematostella*. We generated transgenic reporter lines to show that *NvInsm1*-expressing cells give rise to a large number of sensory and ganglion cells, but not to cnidocytes. We further show that *NvInsm1*::GFP-positive cells co-express *NvTrypsin, NvInsulin-like peptide* (*NvIlp*) and *NvMucin* in different populations of gland/secretory cells in the pharyngeal and body wall ectoderm. Using double transgenics, we observe that *NvSoxB(2)*-expressing cells give rise not only to neurons, but also to *NvInsm1*::GFP-positive gland/secretory cells, and we find that knockdown of *NvSoxB(2)* results in the downregulation of *NvInsm1* and gland/secretory cell genes. Our data show that neurons and gland/secretory cells in *Nematostella* originate from a common pool of progenitor cells and that their development involves *NvInsm1*. These findings support the hypothesis that neurons and secretory cells share a common evolutionary origin.

## RESULTS

### *NvInsm1* is expressed in non-proliferating, differentiating neural cells

The *Nematostella* genome encodes a single Insulinoma-associated protein 1 ortholog (*NvInsm1*) with six C2H2 Zn-finger domains and a SNAG domain at the N-terminus **(Figure S1A**). The domain composition of NvINSM1 is more similar to mouse INSM1 (five Zn-fingers and a SNAG domain) than the *D. melanogaster* and *C. elegans* orthologs, which each have three Zn-fingers and lack the SNAG domain **(Figure S1A, B)**, that is required for interaction with chromatin modifying proteins (Monaghan et al., 2017; Stivers et al., 2000; Welcker et al., 2013). Phylogenetic analysis based on the first three Zn-finger domains confirmed the assignment of NvInsm1 as an Insm1 ortholog **(Figure S1C)**. By *in situ* hybridization, the expression of *NvInsm1* starts at mid-blastula stage in few scattered cells in the epithelium **(Figure 1B, C)**. At late blastula and early gastrula stages, the number of *NvInsm1*-expressing cells increases and is restricted to the aboral half of the embryo (**Figure 1D, E, I**). At late gastrula stage, cells at the oral side also initiate expression **(Figure 1F, J)**. *NvInsm1* starts being expressed in many pharyngeal cells at early planula stage **(Figure 1G, K)** and by late planula stage, it can be detected in mesendodermal single cells **(Figure 1H, L)**. To test whether *NvInsm1* is expressed in proliferating cells, we exposed wild type animals at planula stage to the thymidine analogue EdU, fixed and performed EdU detection together with immunostaining using an antibody raised against NvINSM1 (**Figure S2J, Figure 1M-P)**. After a 30min EdU pulse we could not detect cells expressing NvINSM1 and incorporating EdU. After 2h and 4h of EdU pulse, we could observe a very small number of NvINSM1^+^ cells incorporating EdU (0.56% and 2.27%, respectively) **(Figure 1O, P**). EdU pulse-labelling in combination with fluorescence in situ hybridization likewise did not reveal proliferating *NvInsm1*-expressing cells **(Figure S3)**. This suggests that *NvInsm1* is mostly expressed in non-proliferating cells.

To characterize the nature of the *NvInsm1* expressing cells further, we generated stable transgenic reporter lines, *NvInsm1*::GFP and *NvInsm1*::mCherry, in which a 4,6kb region upstream of the open reading frame of *NvInsm1* drives the expression of a membrane-tethered GFP (or membrane-tethered mCherry). This allows the identification of the *NvInsm1* expressing cells and their progeny during development. Double fluorescent *in situ* hybridization of *GFP* and *NvInsm1* on transgenic embryos showed a high degree of co-expression of the reporter gene transcripts and endogenous *NvInsm1* **(Figure S4)**, confirming that the reporter lines reflect the endogenous expression of *NvInsm1*.

Analysis of the *NvInsm1*::*GFP* line reveals that the transgene is expressed from early gastrula stage in scattered ectodermal cells. At this stage, the cells are elongated along the apical-basal axis, with some having a cilium and others microvilli-like protrusions on their apical surface (**Figure 2A-B**). At planula stage, many of the GFP^+^ ectodermal cells form neurites with varicosities on their basal side (**Figure 2C-D**). At primary polyp stage, GFP labels the ectodermal and mesendodermal nerve net of the animal (**Figure 2E-G**). At this stage there is also GFP expression in many cells of the pharynx and in the ectodermal parts of the two primary mesenteries (the septal filaments, **Figure 2H-M**). Most GFP^+^ cells in the pharynx are elongated along their apical-basal axis and have a cilium on the apical membrane (**Figure 2I**), while some cells are not elongated and seem to contain many vesicle-like structures labelled by the membrane-tethered GFP (**Figure 2J**). Among the vesicle-filled cells, we observe some that also carry an apical cilium (**Figure 2M**). The *NvInsm1::GFP* reporter line thus labels cells with different morphologies, including cells of neuronal appearance.

**Figure 2:**
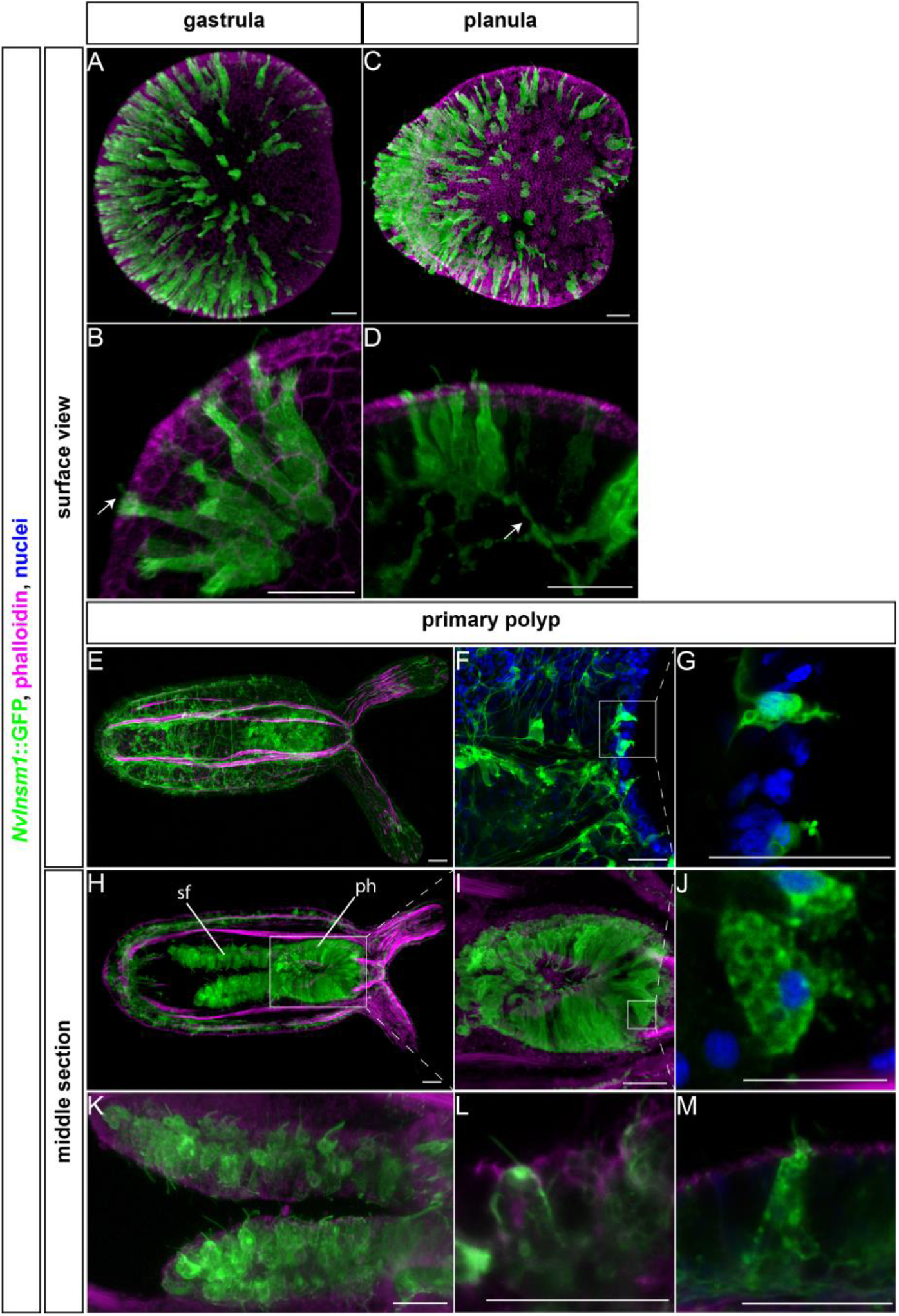
A *NvInsm1* transgenic reporter line labels neurons and gland/secretory cells. (A-M) Confocal microscopy images of the *NvInsm1*::GFP transgenic line, GFP is detected by anti-GFP antibody (green), phalloidin is labeled (magenta) and DAPI (blue), except (F. G) which show live imaging of GFP (green) and Hoechst (blue). Developmental stages are indicated on top and the middle or surface view indicated on the left side. All images are lateral views with the aboral pole to the left. The white arrow in (B) point towards a cell surface with microvilli. The white arrow in (D) shows the forming neurites. (I) is a higher magnification of the area indicated in (H), (G) and (J) are higher magnifications of the boxed areas in (F) and (J), respectively. All images are Imaris snapshots from 3D reconstructions, except images G, J, L and M which are stacks (two to three confocal sections) generated with Fiji. Pictures F and G were taken from live animals (see Material and Methods). Expression of GFP is detected from gastrula stage on and is consistent with the *in situ* hybridization signals (**Figure S2**). ph, pharynx; sf, septal filament. Scale bar represents 20 μm

### *NvInsm1*-expressing cells give rise to sensory and ganglion neurons

To better characterize the identity of the *NvInsm1*-expressing cells, we used double fluorescent *in situ* hybridization to test co-expression of *NvInsm1* with the neuropeptide gene *NvRFamide* (labelling a subset of differentiating sensory and ganglion cells, (Marlow et al., 2009), **Figure 3A-H**). We observed that *NvInsm1* and *NvRFamide* partially co-express in the ectoderm from blastula to planula stage **(Figure 3A-F)** and in the developing pharynx at planula stage (**Figure 3G, H**).

**Figure 3:**
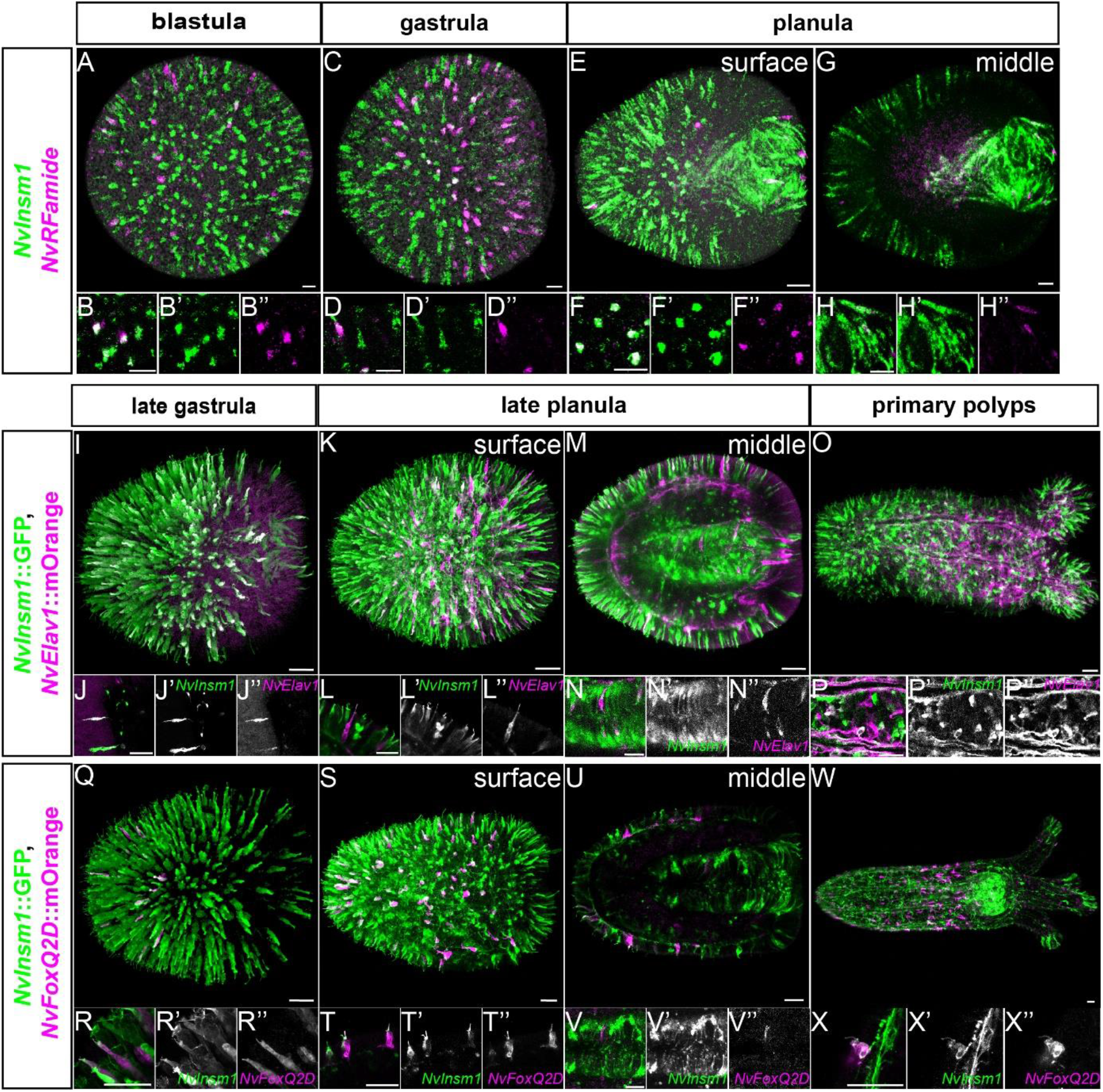
*NvInsm1* identifies sensory and ganglion cells. Developmental stages are indicated at the top and the various labeling on the left. (A-H) Double fluorescence *in situ* hybridization for *NvInsm1* (green) and the neuronal differentiating marker *NvRFamide* (magenta) shows co-expression from blastula stage to planula stage. (B, D, F) are higher magnifications of ectodermal regions at blastula, gastrula and planula, respectively. (H) is a higher magnification of the forming pharynx. (I-P) double transgenic animals with *NvInsm1*::GFP (green) and *NvElav1*::mOrange (magenta). Almost all *NvElav1*::mOrange^+^ cells are GFP^+^. (Q-X) Double transgenic animals with *NvInsm1*::GFP (green) and *NvFoxQ2d*::mOrange (magenta). From gastrula to primary polyps most of the *NvFoxQ2d*::mOrange^+^ cells are also GFP^+^. (J, L, R, T and X) show higher magnifications of ectodermal cells, (N and M) identifies co-expression in the pharynx, (P) shows co-expression in mesendodermal neurons in the body column. All pictures are lateral views with aboral pole to the left. (A, C, E, I, K, O, Q, S and W) are Imaris snapshots from 3D reconstructions to show the overview. All other pictures are stacks of two to three confocal sections to ensure the co-expression of the signals. Scale bars correspond to 20 μm

Next, we generated double transgenic animals by crossing *NvInsm1*::GFP animals with previously characterized transgenic reporter lines. The *NvElav1*::mOrange transgenic line labels a large fraction of sensory and ganglion cells but not cnidocytes (Nakanishi et al., 2012). At late gastrula, all *NvElav1*::mOrange^+^ cells are also *NvInsm1*::GFP^+^ suggesting that a subset of the *NvInsm1*-expressing cells are sensory and ganglion cells (**Figure 3I-J**). From late planula to primary polyp stage most of the *NvElav1*::mOrange^+^ cells are still expressing the GFP transgene, including cells present in the pharynx and in the forming tentacles (**Figure 3K-P**). The *NvFoxQ2d*::mOrange line labels a small population of ectodermal sensory cells which does not co-express *NvElav1::*cerulean (Busengdal and Rentzsch, 2017). In double transgenics we identified the *NvFoxQ2d::*mOrange^+^ cells as a subpopulation of the *NvInsm1*::GFP^+^ cells (**Figure 3Q-X**). These observations were also confirmed by studying the presence of NvINSM1 protein in those transgenic lines (**Figure S2A-I**).

Thus, the *NvInsm1* transgenic reporter line identifies two of the main classes of neurons in *Nematostella*, sensory cells and ganglion cells.

### *NvInsm1* expressing cells do not give rise to cnidocytes

We next analysed whether *NvInsm1*-expressing cells also give rise to the third class of cnidarian neural cells, the cnidocytes. First, we studied *NvInsm1* expression with *NvNcol3* (a gene expressed in differentiating cnidocytes) via double fluorescent *in situ* hybridization (**Figure S5A-H)**. The two genes do not colocalize from blastula to planula stage, but we cannot discard the possibility that they are expressed sequentially in the same cells. In order to investigate this hypothesis, we generated double transgenics with the *NvNcol3*::mOrange2 line, which identifies differentiating and mature cnidocytes (Sunagar et al., 2018). The two transgenes do not co-express from the onset of the *NvNcol3*::mOrange2 expression at late gastrula stage to primary polyp stage (**Figure S5I-P**). Based on these two analyses we conclude that *NvInsm1-*expressing cells do not contribute to the formation of cnidocytes.

### Some *NvInsm1* expressing cells develop into gland/secretory cells

From planula to primary polyp stages, the number of *NvInsm1-*expressing cells in the developing pharynx and in the septal filaments along the mesenteries seemed to increase. These two structures contain many neural cells, but they have also been described to contain many gland/secretory cells expressing genes encoding digestive enzymes or genes related to mucus secretion (Steinmetz et al., 2017). To test whether *NvInsm1*-expressing cells contribute to populations of gland/secretory cells, we performed fluorescence *in situ* hybridization in the background of the *NvInsm1::GFP* line at planula stage (**Figure 4A-F)**. We observed that *NvInsm1*::GFP is co-expressed with *NvMucin* in putative mucous cells of the ectoderm of the body column and in the forming pharynx (**Figure 4A, B** (Steinmetz et al., 2017). We also observed co-expression in the pharynx with *NvTrypsinA* (expressed in putative gland cells with digestive function, **Figure 4C, D)** as well as with Insulin-like peptide 2 (*NvILP2*; **Figure 4E, F)** (Steinmetz et al., 2017). We conclude that a subpopulation of the *NvInsm1*-expressing cells gives rise to different types of gland/secretory cells in *Nematostella*.

**Figure 4.**
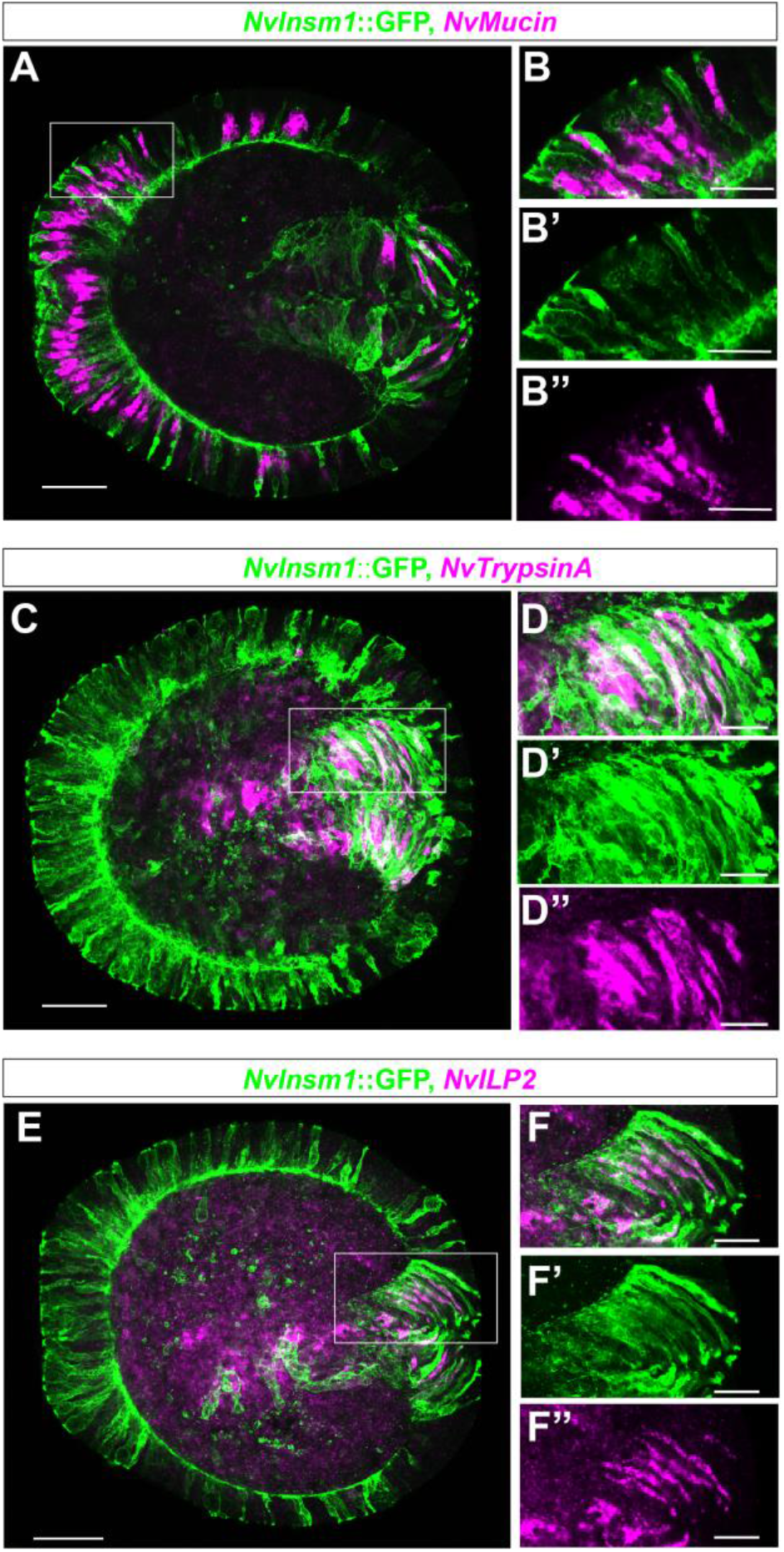
The *NvInsm1*::GFP reporter line labels cells expressing markers for gland/secretory cells. (A-F) Confocal images of fluorescence *in situ* hybridizations in *NvInsm1*::GFP transgenic planulae. GFP (green) is detected by immunohistochemistry, in situ probes are indicated on the top and shown in magenta. (B, D and F) are higher magnifications of the boxed areas in (A, C and E), respectively. Mid-lateral views with the aboral pole to the left. Transcripts for *NvMucin, NvTrypsinA* and *NvILP2* are detected in *NvInsm1*::GFP positive cells. All pictures are stacks of 1-3 confocal sections. Scale bars represent 50μm (A, C and E) and 20 μm (B, D, F).

The published *Nematostella* single cell sequencing data confirm our morphological observation by identifying *NvInsm1* in neuronal and in gland/secretory metacells (Sebe-Pedros et al., 2018). To understand whether this expression profile is conserved in other cnidarians, we used the single cell RNA sequencing analysis performed in *Hydra* (Siebert et al., 2019), a distantly related cnidarian species. The *Hydra* genome encodes a single *Insm1-like* gene that contains two C2H2 zinc fingers (and was therefore not included in our phylogenetic analysis) that show strong similarity to Insm1 proteins from other species. *HvInsm1-like* appears to be present in ectodermal and endodermal neurons as well as in gland/secretory cells and in their progenitors (**Figure S6**). As in *Nematostella, HvInsm1-like* is not expressed in differentiated cnidocytes and in nematoblasts (cnidocyte progenitors). Altogether, these data suggest that in cnidarians *Insm1* is expressed in sensory and ganglion cells as well as in gland/secretory cells.

### *NvSoxB(2)*-expressing progenitor cell population give rise to neural and gland cells

During *Nematostella* embryogenesis, a *NvSoxB(2)*-expressing population of cells gives rise to sensory cells, ganglion cells and cnidocytes (Richards and Rentzsch, 2014). Comparison of in situ hybridization at different developmental stages (**Figure S7A-N**) showed that the expression of *NvSoxB(2)* in the blastoderm commences earlier than that of *NvInsm1* (**Figure S7A, H**) and that, similarly, expression of *NvSoxB(2)* in the pharynx and the mesendoderm precedes that of *NvInsm1* (**Figure S7E, F, L, M**). In double fluorescence in situ hybridizations (**Figure 5 A-H**), we observed many cells co-expressing the two transcripts at blastula, gastrula and planula stages, but we also found cells expressing only one of the two genes. This is compatible with at least two, not mutually exclusive scenarios: individual cells might first express *NvSoxB(2)*, then co-express *NvSoxB(2)* and *NvInsm1*, and eventually maintain only the expression of *NvInsm1*. Alternatively, the double-positive and single-positive cells might constitute different populations with different developmental potentials. To test these hypotheses, we generated *NvSoxB(2)*::mOrange, *NvInsm1*::GFP double transgenic animals (**Figure 5I-Z**). At gastrula, planula and primary polyp stage, all detectable *NvInsm1*::GFP^*+*^ cells were also labelled with the *NvSoxB(2)* reporter (**Figure 5J, M, N, P, S and T)**, but there were some *NvSoxB(2)::*mOrange^+^ cells that were not GFP^+^, suggesting that the developmental potential of *NvSoxB(2)* expressing cells is broader than that of the *NvInsm1*^*+*^ ones (**Figure 5K, Q**). If all *NvInsm1::*GFP^+^ cells are part of the *NvSoxB(2)* lineage, we would expect to find gland/secretory cells among the *NvSoxB(2)*::mOrange cells. This was indeed the case, as we identified double transgenic ectodermal cells containing vesicle-like structures (**Figure 5X, Y, Z**). This suggest that *NvSoxB(2)*-expressing cells constitute a population of progenitor cells that gives rise to neural cells and gland/secretory cells.

**Figure 5:**
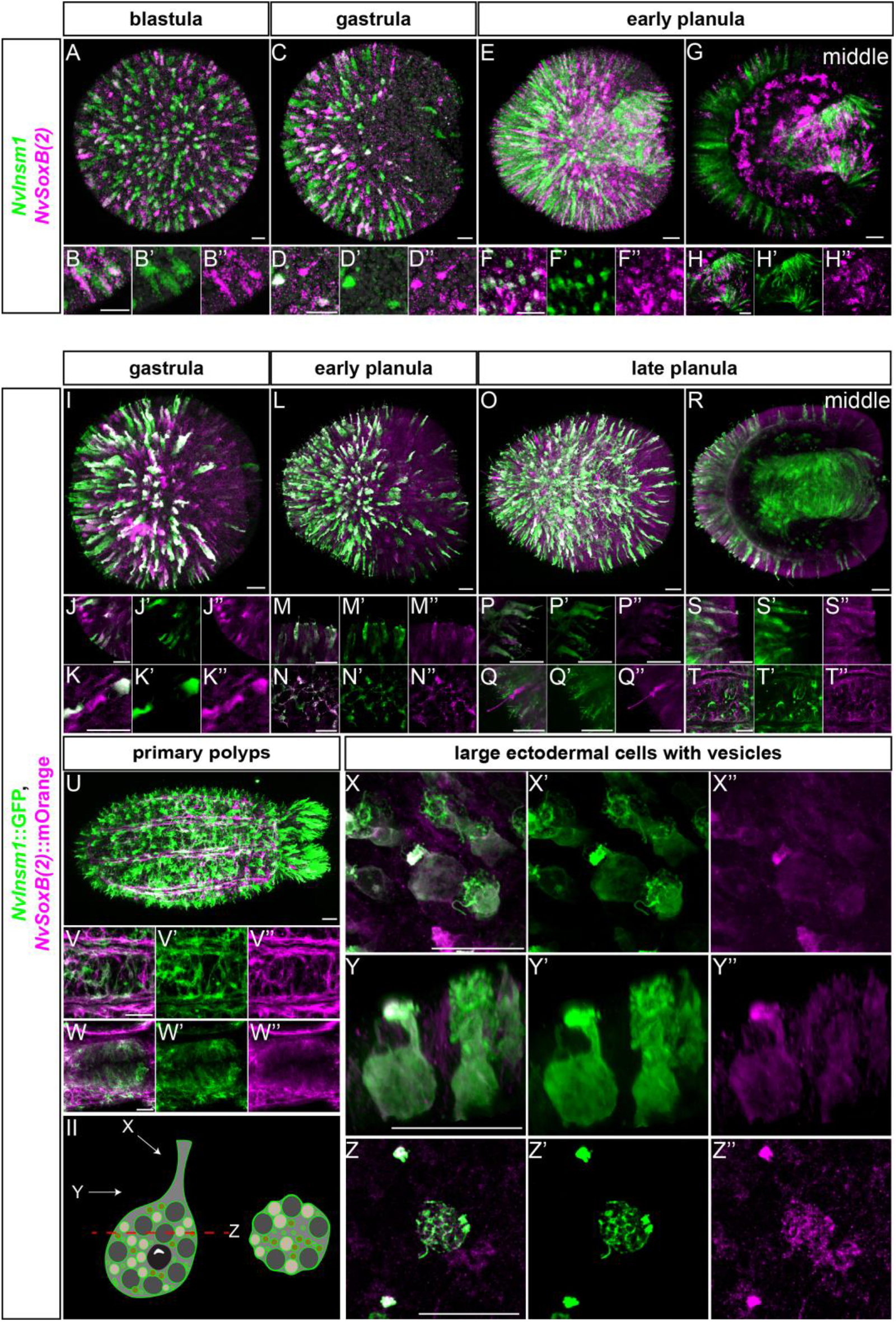
*NvInsm1* identifies a subset of the *NvSoxB(2)* expressing cell progeny. For (A-Z), probes or transgenes are represented on the left and stages at the top. (A-H) Double fluorescence *in situ* hybridization at blastula stage for *NvInsm1* (green) and *NvSoxB(2)* (magenta) shows co-expression from blastula to early planula stage in ectodermal cells (A-F) and in the pharyngeal cells at early planula (G, H). All *in situ* hybridizations were performed with at least three replicates. Overview images A, C and E are Imaris snapshots from 3D reconstructions. All other pictures are Fiji stacks of two confocal sections to ensure the co-expression of the signals. (I-Z) Double transgenic animals with *NvInsm1*::GFP (green) and *NvSoxB(2)*:: mOrange (magenta). Lateral view with aboral pole to the left. From gastrula to primary polyp stage, all *NvInsm1*::GFP^+^ cells are also mOrange^+^. (J, M, S, T) show snapshots of cells at different stages expressing both transgenes, forming neurites are visible in (N). (V) Snapshot of the developing mesendo-dermal nerve net. (W) Snapshot of the developing pharynx, at primary polyp stage. At every stage, some large cells containing many vesicle-like structures also express both transgenes (X, Y, Z). (II) Diagram representation of one of these cells with the different views taken in the (X-Z). (X) Surface view of these cells from a slightly oblique angle. (Y) Lateral view, 3D reconstruction of one cell. (Z) Single confocal cross section of one of the cells in the plane of the epithelium shows the vesicle-like structures. Some *NvSoxB(2)*::mOrange^+^ cells are not GFP^+^ (K, Q), suggesting that *NvInsm1* identifies a subset of the *NvSoxB(2)*-expressing cell population. (I, L, O, U, X and Y) are Imaris snapshots from the 3D reconstruction to show the overview staining. All other pictures (except Z) are Fiji stacks of two to three confocal sections to ensure the co-expression of the signals. Scale bar represents 20 μm.

To test whether *NvSoxB(2)* function is required for the expression of *NvInsm1* and for the development of gland cells, we inhibited *NvSoxB(2)* by injection of a morpholino antisense oligonucleotide (Richards and Rentzsch, 2014). This resulted in a nearly complete suppression of *NvInsm1* expression (**Figure 6A, B**) and a strong reduction in the number of *NvInsm1*::mCherry expressing cells (**Figure 6C-F**). Furthermore, the number of cells expressing *NvMucin* (in pharynx and body wall ectoderm) and *NvTrypsinA* (in the pharynx) was strongly reduced upon injection of *NvSoxB(2)* morpholinos (**Figure 6G-J**). Expression of *NvFoxA* showed that the effect on pharyngeal cells is not due to a general effect on pharynx development (**Figure 6K, L**).

**Figure 6:**
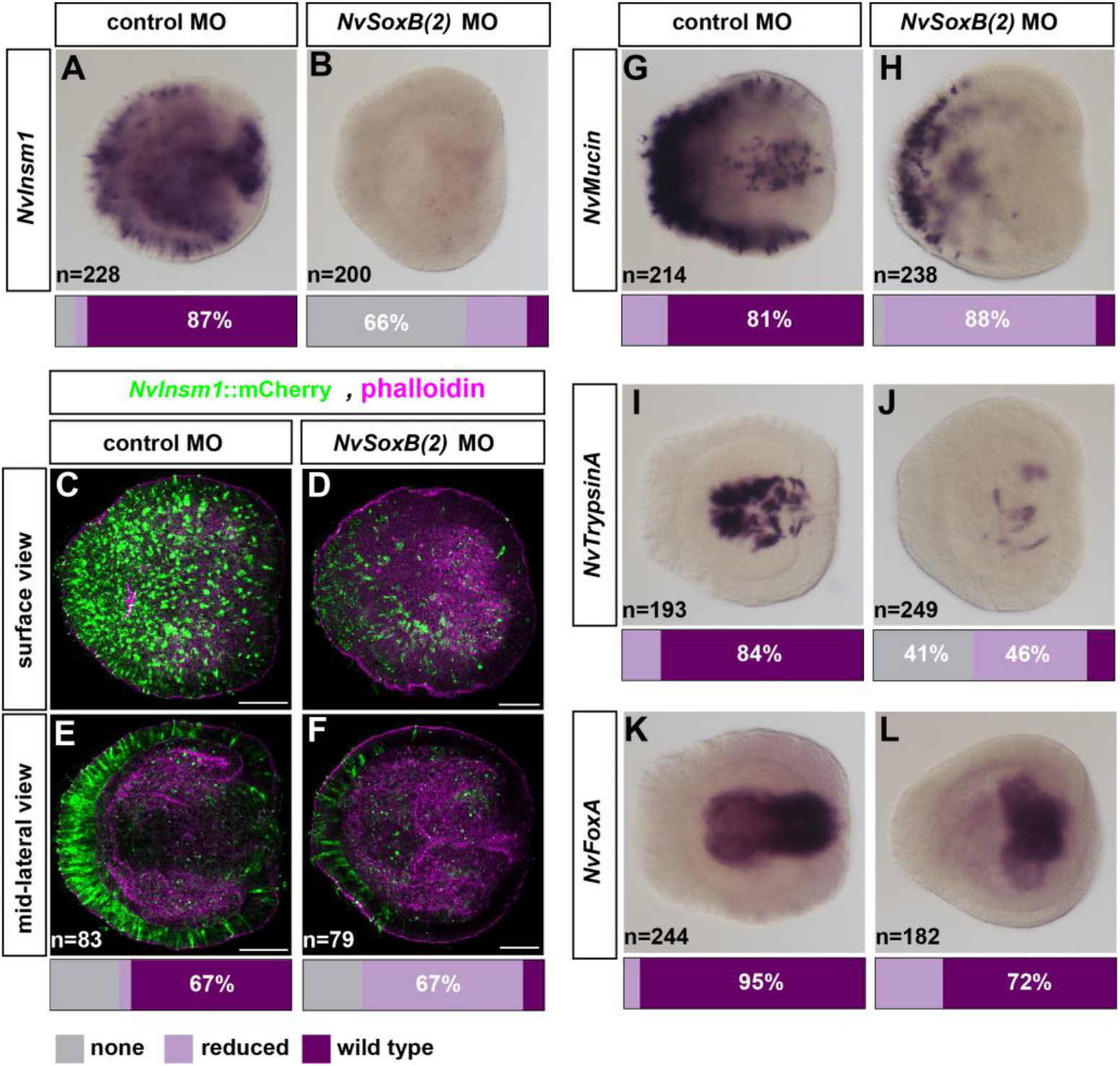
*NvSoxB(2)* is required for the development of gland/secretory cells. (A, B, G-L) Lateral views of *in situ* hybridizations at planula stage with the aboral pole to the left. Injected morpholinos are indicated at the top, probes are indicated on the left side. *NvSoxB(2)* MO injection results in a decreased number of *NvInsm1* (A, B), *NvMucin* (G, H) and *NvTryspinA*-expressing cells (I, J). Expression of *NvFoxA* in the pharynx is unaffected, though the shape of the pharynx is altered (K, L). Animals were quantified into phenotypic classes based on having no, reduced or wild-type expression. Bars at the base of each image represent the percentage of animals in each phenotypic class. (C-F) show a decrease of *NvInsm1*::mCherry^+^ cells (green) after injection of the *NvSoxB(2)* morpholino. Phalloidin staining (magenta) allow the visualization of the morphology of the developing pharynx. Scale bar represents 50 μm.

Taken together, these observations suggest that neural as well as gland cells in *Nematostella* derive from a common *NvSoxB(2)*-expressing progenitor cell population.

## DISCUSSION

In this study we have shown that neurons and gland/secretory cells in *Nematostella* originate from a common population of progenitor cells that is characterized by the expression of *NvSoxB(2)*. Among the cells derived from this progenitor population, *NvInsm1* is expressed in sensory cells, ganglion cells and gland/secretory cells. We propose a scenario in which *NvInsm1* functions at an early stage in the postmitotic differentiation program of these cells and we interpret our findings as support for a close evolutionary relationship of neurons and gland/secretory cells.

We have previously characterized *NvSoxB(2)*-expressing cells as a population that during embryonic development includes proliferating cells (determined by EdU incorporation in pulse-labelling experiments) and that gives rise to the three classes of cnidarian neural cells: sensory cells, ganglion cells and cnidocytes (Richards and Rentzsch, 2014). Focusing on later developmental stages and using double transgenic animals with the brighter *NvInsm1::GFP* line, we find here that *NvSoxB(2)*^+^ cells in addition give rise to different types of gland/secretory cells. Moreover, knockdown of *NvSoxB(2)* results in a strong reduction in the expression of genes expressed in gland/secretory cells. While these observations show that the population of *NvSoxB(2)*-expressing cells has a broader developmental potential than previously thought (see also (Cole et al., 2020), the degree of heterogeneity of progenitor cells in this population remains unknown. It might include progenitors with a developmental potential restricted to one class of cells (e.g. sensory cells or gland/secretory cells) and/or progenitors that can give rise to different classes of cells. These scenarios are not mutually exclusive, as *NvSoxB(2)* expression might be maintained while progenitors undergo divisions and experience a stepwise restriction of their developmental potential. We found that *NvInsm1* and *NvSoxB(2)* transcripts are present simultaneously in many cells during development, but that there are also cells that express only one of the transcripts. In the transgenic reporter line, all *NvInsm1*::GFP expressing cells are part of the *NvSoxB(2)*::mOrange population. Since we did not observe EdU incorporation in *NvInsm1* expressing cells (after a 30min pulse), this suggests that *NvInsm1* transcription commences in cells that already express *NvSoxB(2)*, but do not proliferate any longer and that it is maintained after *NvSoxB(2)* transcription ceases. While *NvInsm1* is thus expressed early in postmitotic precursors of neurons and gland/secretory cells (but not in precursors of cnidocytes), it does not identify proliferating progenitors dedicated to the generation of these two large populations of cells. In *Hydra*, trajectory analyses of single cell RNA sequencing (scRNA seq) data showed that expression of *Hydra Insm1-like* is found in neurons, in gland/secretory cells and in their precursors (Siebert et al., 2019). This suggests that a role for *Insm1* in the development of these cells is common to different groups of cnidarians.

Based on the calcium-dependent ultrafast exocytosis of their extrusive organelle, the presence of synaptic connections and shared developmental gene expression, cnidocytes are traditionally considered a derived neural cell type (Galliot et al., 2009; Nuchter et al., 2006; Oliver et al., 2008; Rentzsch et al., 2017; Westfall, 1996) and we were therefore surprised that *NvInsm1* is not expressed in cnidocytes. In *Hydra*, expression of *Insm1-like* has also not been detected in cnidocytes or their precursors (Siebert et al., 2019). These observations prompt more detailed lineage tracing and functional analyses to obtain a better understanding of the developmental relationship between neurons, cnidocytes and gland cells.

In vertebrates, *Insm1* regulates the formation of cells with secretory capacity but very different developmental origins, e.g. neurons derived from the ectoderm and intestinal endocrine cells derived from the endoderm (Farkas et al., 2008; Gierl et al., 2006; Horb et al., 2009; Jacob et al., 2009; Wildner et al., 2008). In *Nematostella*, neurons are generated in ectoderm and mesendoderm (Nakanishi et al., 2012), whereas gland cells expressing digestive enzymes are generated in the pharyngeal ectoderm and in the septal filaments of the mesenteries, a derivative of the pharyngeal ectoderm (Steinmetz et al., 2017). As in vertebrates, *NvInsm1* is thus expressed in cells in which tightly regulated secretion is thought to be of importance, but which have different embryological origins. One possible explanation for this observation would be a function in the regulation of genes directly involved in the secretion process, e.g. in the biogenesis, transport or release of secretory vesicles. This scenario is, however, not well supported by data from other model organisms. In mice, *Insm1* regulates neurogenesis in the cortex, the olfactory epithelium and the otocyst by controlling the development of proliferative progenitors (Farkas et al., 2008; Lorenzen et al., 2015; Rosenbaum et al., 2011; Tavano et al., 2018). In the development of lung and pituitary endocrine cells, of serotonergic and adrenergic neurons, and of zebrafish motor neurons, *Insm1* instead is required for terminal differentiation (Gong et al., 2017; Jacob et al., 2009; Jia et al., 2015a; Welcker et al., 2013) and this can include the repression of alternative differentiation programs (Welcker et al., 2013; Wiwatpanit et al., 2018). Despite the striking restriction of *Insm1* function to neural and endocrine cells, there are thus likely differences in the specific regulatory programs regulated by *Insm1* in different populations of cells in vertebrates. Comparisons of Insm1 function across a broader range of animals and identification of direct target genes will likely be informative for obtaining a better understanding of the evolution of Insm1 function.

In contrast to the *Drosophila* and *C. elegans Insm1* orthologs, *NvInsm1* encodes an N-terminal SNAG motif that in mice mediates interaction with chromatin modifying complexes containing Lsd1/KDM1A, CoREST and HDAC1/2 (Monaghan et al., 2017; Welcker et al., 2013). The SNAG motif has been shown to be essential for the function of mouse Insm1 in pituitary development (Welcker et al., 2013) and the presence of this motif in *Nematostella Insm1* might facilitate cross-species comparisons of the transcriptional regulation by Insm1 genes.

Our observations reveal developmental commonalities of cnidarian neurons and gland/secretory cells at the level of cell lineage and gene regulation. Together with previously described similarities in the development of these cell types among bilaterian species, our findings support the hypothesis that neurons and endocrine cells of extant animals evolved from primordial sensory-secretory cells (Grundfest, 1965; Hartenstein et al., 2017; Jekely, 2021).

## MATERIALS AND METHODS

### *Nematostella* culture

Animals were maintained at 18°C in 1/3 filtered seawater (=*Nematostella* medium, NM). Spawning induction was performed by light and temperature shift as described in (Fritzenwanker and Technau, 2002). Incubation of the fertilized egg packages with 3% cysteine/NM (pH 7.4) removed the jelly. Embryos were then raised at 21 °C and fixed at 12hpf (early blastula), 16hpf (blastula), 20hpf (early gastrula), 24hpf (gastrula), 30hpf (late gastrula), 48hpf (early-planula), 72hpf (planula), 4dpf (late planula); 5dpf (tentacle bud); 7dpf (primary polyp).

### Identification of *NvInsm1* and generation of transgenic lines

*NvInsm1* is derived from gene model Nve7206, retrieved from https://figshare.com/articles/Nematostella_vectensis_transcriptome_and_gene_models_v2_0/807696. The *NvInsm1*::GFP transgenic reporter line was generated by meganuclease mediated transgenesis as described by (Renfer et al., 2009; Renfer and Technau, 2017). The genomic coordinates for the 4,6 kb regulatory region are 179.529-184.145 on scaffold 185 (http://genome.jgi.doe.gov/Nemve1/Nemve1.home.html, accessed 15 August 2019). This fragment was inserted in front of a codon optimized GFP via the HiFi DNA Assembly kit (NEB) with the addition of a membrane-tethering CAAX domain at the C-terminus to visualize the morphology of the cells expressing the reporter protein. The *NvSoxB(2)*::mOrange line has been described in (Richards and Rentzsch, 2014), *NvElav1*::mOrange in (Nakanishi et al., 2012), *NvNCol3*::mOrange2 in (Sunagar et al., 2018) and *NvFoxQ2d*::mOrange in (Busengdal and Rentzsch, 2017).

### *In situ* hybridization, EdU labelling and immunohistochemistry

In situ hybridization (ISH) and fluorescence ISH (FISH) were performed as described in the Supplementary material in (Richards and Rentzsch, 2014). For combination of FISH and immunohistochemistry (IHC), FISH was performed first using the TSA plus Cyanine 3 kit (Perkin Elmer NEL744001KT), followed by IHC with anti-GFP primary antibody (abcam290) and goat anti-rabbit Alexa 488 secondary antibody (Invitrogen A11011).

Other primary antibodies were: to detect *NvInsm1*::GFP (without FISH), anti-GFP (mouse, abcam1218, 1:200); anti-dsRed, to detect mOrange (rabbit, Clontech 632496, 1:100). The polyclonal NvINSM1 antibody was raised by GenScript in rabbit against amino acids 3 – 170 of NvINSM1 expressed in and purified from *E*.*coli*. The affinity-purified antibody was used at 1:100 dilution, incubation lasted for 48h. EdU labelling was done as 30min, 2h and 4h pulses followed by fixation as described in (Richards and Rentzsch 2014), using Click-it EdU Alexa fluor 488 and 647 kits (Molecular Probes C10337). For quantification, a square of 100 x 100µm in the body column (see Figure 1L) was analyzed after imaging by confocal microscopy.

### Microscopy and image processing

Samples were imaged either on a Nikon Eclipse E800 compound microscope with a Nikon Digital Sight DSU3 camera (for colorimetric in situ hybridizations) or on a Leica SP5 confocal microscope. For Figure 4, samples were imaged on an Olympus Fluoview FV3000 confocal microscope. Live imaging was performed on primary polyps (between 10-14 dpf). Polyps were live stained with Hoechst 33342 (ThermoFisher Scientific 62249) at 1:2000 in NM for 10min at room temperature. Animals were then relaxed by slowly adding 1M MgCl_2_ and incubating the fully extended animals for 10 min. They were then mounted in this solution on a glass slide and imaged at the confocal microscope. Image manipulation was performed with Imaris (3D reconstructions of whole stacks to present embryos overviews) and with Fiji for zoomed snapshots (single picture or stack of 2-3 confocal sections). Figures were assembled in Adobe Illustrator.

### Western blotting

Protein extraction was performed on wild-type late planula. Animals were placed in RIPA buffer (150mM NaCl, 50mM Tris pH8, 1% NP40, 0.5% DOC, 0.1% SDS) supplemented with complete EDTA-free Protease Inhibitor Cocktail (Roche, 4693159001) and homogenized by passing through a 27G needle. Samples were incubated on ice for 30 minutes and mixed by passing through the needle every 5 minutes and centrifuged at full speed for 15 minutes @ 4°C. The supernatant was kept and the protein concentration quantified using the Qubit™ Protein Assay (Invitrogen, Q33212). 30 μg of protein was used per lane, mixed 1:1 with 2X Laemmli sample buffer (0.1M TrisHCl pH 6.8, 2% SDS, 20% Glycerol, 4% βmercaptoethanol 0.02% Bromophenol blue) and boiled for 5 minutes before loading. PageRuler™ Plus prestained protein ladder, 10 to 250 kDa (Thermo Scientific, 26619) was used. SDS PAGE was performed using 7.5% Mini-PROTEAN® TGX™ precast protein gels (BIO-RAD, 4561023) run in running buffer (25mM Tris, 192mM Glycine, 0.1% SDS) at 100 V for ∼120 minutes. Transfer was performed using Trans-Blot Turbo Mini 0.2 µm PVDF Transfer Pack (BIO-RAD, 1704156) on a Trans-Blot Turbo transfer system (BIO-RAD) using the high molecular weight program. After transfer the membrane was washed in PBT (PBS + 0.1% Tween) several times and blocked with 5% milk powder in PBT (MPBT) at RT for 1 hour. The blots were incubated o/n at 4°C in primary antibody (Rabbit Anti-NvInsm1, concentration 1:10000) in MPBT. The membranes were then washed several times in PBT and incubated in secondary antibody (Goat anti-Rabbit (HRP), Ab97051, concentration 1:10000) in MPBT at RT for 1 hour. Membranes were then washed several times in TBT and the signal was revealed using Clarify ECL substrate (BIO-RAD, 1705060) and imaged on a ChemiDoc XRS+ (BIO-RAD). The blots were then washed in PBT and blocked again in 5% MPBT for 1 hour at RT.

### Morpholino injection

*NvSoxB(2)* MO and the generic control MO are as in (Richards and Rentzsch, 2014). Experiments were conducted with three biological replicates, with embryos derived from three independent spawnings. Morpholino sequences are: *NvSoxB(2)* MO1: TATACTC TCCGCTGTGTCGCTATGT, control MO: CCATTGTGAAGTTAAACGATAGATC

### Phylogenetic analysis

Amino acid sequences (Table S1) were retrieved by BLAST searches at NCBI and via OrthoDB (https://www.orthodb.org/) (Kriventseva et al., 2019). Zinc finger domains were determined by SMART (Letunic et al., 2021), only the first three zinc fingers were used for the alignment. Sequences before zinc finger 1 were removed, and so were the sequences between zinc fingers 2 and 3. *Hydra, Trichoplax* and *Clytia* genes with best reciprocal Blast hits to Insm1 were omitted because they only have two zinc fingers. *Snail* sequences from *Nematostella vectensis* and *Homo sapiens* were used as outgroup. The alignment was generated with Muscle 3.7 (Edgar, 2004) at Phylemon 2 (http://phylemon2.bioinfo.cipf.es). The phylogenetic analysis was run with PhyML3.1 (Guindon and Gascuel, 2003) and the WAG model at www.phylogeny.fr (Dereeper et al., 2008) with 100 bootstraps.

### Analysis of Hydra single cell data

The *Hydra* single cell analysis (Siebert et al., 2019) give access to the Broad Single-Cell Portal available at: https://portals.broadinstitute.org/single_cell/study/SCP260/stem-cell-differentiation-trajectories-in-hydra-resolved-at-single-cell-resolution. On this portal, genes can be analysed graphically and each graphical picture can be saved.

## ACKNOWLEDGEMENTS

We thank Eilen Myrvold and Lavina Jubek for excellent care of the *Nematostella* facility. Pawel Burkhardt and Tarja Hoffmeyer for sharing reagents and providing advice on western blotting, Johanna Kraus for advice on live imaging, Marios Chatzigeorgiou for helpful discussions and advice and Gemma Richards for initial in situ hybridizations. The work was funded by the Sars Centre Core budget and by a grant from the Research Council of Norway and the University of Bergen (251185/F20) to F.R..

## REFERENCES

Apelqvist, A., Li, H., Sommer, L., Beatus, P., Anderson, D. J., Honjo, T., Hrabe de Angelis, M., Lendahl, U. and Edlund, H. (1999). Notch signalling controls pancreatic cell differentiation. Nature 400, 877–881.

Arendt, D. (2020). The Evolutionary Assembly of Neuronal Machinery. Curr Biol 30, R603–R616.

Arntfield, M. E. and van der Kooy, D. (2011). beta-Cell evolution: How the pancreas borrowed from the brain: The shared toolbox of genes expressed by neural and pancreatic endocrine cells may reflect their evolutionary relationship. BioEssays 33, 582–587.

Bosch, T. C., Anton-Erxleben, F., Hemmrich, G. and Khalturin, K. (2010). The Hydra polyp: nothing but an active stem cell community. Dev Growth Differ 52, 15–25.

Breslin, M. B., Zhu, M. and Lan, M. S. (2003). NeuroD1/E47 regulates the E-box element of a novel zinc finger transcription factor, IA-1, in developing nervous system. The Journal of biological chemistry 278, 38991–38997.

Brunet, T. and Arendt, D. (2016). From damage response to action potentials: early evolution of neural and contractile modules in stem eukaryotes. Philosophical transactions of the Royal Society of London. Series B, Biological sciences 371, 20150043.

Buas, M. F. and Kadesch, T. (2010). Regulation of skeletal myogenesis by Notch. Exp Cell Res 316, 3028–3033.

Burkhardt, P. and Sprecher, S. G. (2017). Evolutionary origin of synapses and neurons - Bridging the gap. BioEssays: news and reviews in molecular, cellular and developmental biology 39.

Busengdal, H. and Rentzsch, F. (2017). Unipotent progenitors contribute to the generation of sensory cell types in the nervous system of the cnidarian Nematostella vectensis. Dev Biol 431, 59–68.

Cau, E., Casarosa, S. and Guillemot, F. (2002). Mash1 and Ngn1 control distinct steps of determination and differentiation in the olfactory sensory neuron lineage. Development 129, 1871–1880.

Cole, A. G., Kaul, S., Jahnel, S. M., Steger, J., Zimmerman, B., Reischl, R., Richards, G. S., Rentzsch, F., Steinmetz, P. and Technau, U. (2020). Muscle cell type diversification facilitated by extensive gene duplications. bioRxiv, 2020.2007.2019.210658.

de la Pompa, J.L., Wakeham, A., Correia, K. M., Samper, E., Brown, S., Aguilera, R. J., Nakano, T., Honjo, T., Mak, T. W., Rossant, J., et al. (1997). Conservation of the Notch signalling pathway in mammalian neurogenesis. Development 124, 1139–1148.

Dereeper, A., Guignon, V., Blanc, G., Audic, S., Buffet, S., Chevenet, F., Dufayard, J. F., Guindon, S., Lefort, V., Lescot, M., et al. (2008). Phylogeny.fr: robust phylogenetic analysis for the non-specialist. Nucleic acids research 36, W465–469.

dos Reis, M., Thawornwattana, Y., Angelis, K., Telford, M. J., Donoghue, P. C. and Yang, Z. (2015). Uncertainty in the Timing of Origin of Animals and the Limits of Precision in Molecular Timescales. Curr Biol 25, 2939–2950.

Duggan, A., Madathany, T., De Castro, S. C. P., Gerrelli, D., Guddati, K. and Garcia-Anoveros, J. (2008). Transient expression of the conserved zinc finger gene INSM1 in progenitors and nascent neurons throughout embryonic and adult neurogenesis. Journal of Comparative Neurology 507, 1497–1520.

Dunn, C. W., Giribet, G., Edgecombe, G. D. and Hejnol, A. (2014). Animal Phylogeny and Its Evolutionary Implications. Annu Rev Ecol Evol S 45, 371–395.

Edgar, R. C. (2004). MUSCLE: multiple sequence alignment with high accuracy and high throughput. Nucleic acids research 32, 1792–1797.

Ernsberger, U. (2015). Can the ‘neuron theory’ be complemented by a universal mechanism for generic neuronal differentiation. Cell and Tissue Research 359, 343–384.

Farkas, L. M., Haffner, C., Giger, T., Khaitovich, P., Nowick, K., Birchmeier, C., Paabo, S. and Huttner, W. B. (2008). Insulinoma-associated 1 has a panneurogenic role and promotes the generation and expansion of basal progenitors in the developing mouse neocortex. Neuron 60, 40–55.

Frank, U., Plickert, G. and Muller, W. A. (2009). Cnidarian interstitial cells: The dawn of stem cell research. In Stem Cells in Marine Organisms (ed. B. Rinkevich & V. Matranga), pp. 33–59: Springer.

Fritzenwanker, J. H. and Technau, U. (2002). Induction of gametogenesis in the basal cnidarian Nematostella vectensis(Anthozoa). Dev Genes Evol 212, 99–103.

Galliot, B. and Quiquand, M. (2011). A two-step process in the emergence of neurogenesis. The European Journal of Neuroscience 34, 847–862.

Galliot, B., Quiquand, M., Ghila, L., de Rosa, R., Miljkovic-Licina, M. and Chera, S. (2009). Origins of neurogenesis, a cnidarian view. Dev Biol 332, 2–24.

Gierl, M. S., Karoulias, N., Wende, H., Strehle, M. and Birchmeier, C. (2006). The zinc-finger factor Insm1 (IA-1) is essential for the development of pancreatic beta cells and intestinal endocrine cells. Genes & Development 20, 2465–2478.

Gong, J., Wang, X., Zhu, C., Dong, X., Zhang, Q., Wang, X., Duan, X., Qian, F., Shi, Y., Gao, Y., et al. (2017). Insm1a Regulates Motor Neuron Development in Zebrafish. Front Mol Neurosci 10, 274.

Gradwohl, G., Dierich, A., LeMeur, M. and Guillemot, F. (2000). neurogenin3 is required for the development of the four endocrine cell lineages of the pancreas. Proc Natl Acad Sci U S A 97, 1607–1611.

Grundfest, H. (1965). Evolution of electrophysiological varieties among sensory receptor systems. In Essays on Physiological Evolution (ed. J.W.S. Pringle), pp. 107–138. London: Pergamon Press.

Guindon, S. and Gascuel, O. (2003). A simple, fast, and accurate algorithm to estimate large phylogenies by maximum likelihood. Syst Biol 52, 696–704.

Hartenstein, V. and Stollewerk, A. (2015). The evolution of early neurogenesis. Dev Cell 32, 390–407.

Hartenstein, V., Takashima, S., Hartenstein, P., Asanad, S. and Asanad, K. (2017). bHLH proneural genes as cell fate determinants of entero-endocrine cells, an evolutionarily conserved lineage sharing a common root with sensory neurons. Dev Biol 431, 36–47.

Horb, L. D., Jarkji, Z. H. and Horb, M. E. (2009). Xenopus insm1 is essential for gastrointestinal and pancreatic endocrine cell development. Dev Dyn 238, 2505–2510.

Jacob, J., Storm, R., Castro, D. S., Milton, C., Pla, P., Guillemot, F., Birchmeier, C. and Briscoe, J. (2009). Insm1 (IA-1) is an essential component of the regulatory network that specifies monoaminergic neuronal phenotypes in the vertebrate hindbrain. Development 136, 2477–2485.

Jekely, G. (2021). The chemical brain hypothesis for the origin of nervous systems. Philosophical transactions of the Royal Society of London. Series B, Biological sciences 376, 20190761.

Jenny, M., Uhl, C., Roche, C., Duluc, I., Guillermin, V., Guillemot, F., Jensen, J., Kedinger, M. and Gradwohl, G. (2002). Neurogenin3 is differentially required for endocrine cell fate specification in the intestinal and gastric epithelium. EMBO J 21, 6338–6347.

Jia, S., Ivanov, A., Blasevic, D., Muller, T., Purfurst, B., Sun, W., Chen, W., Poy, M. N., Rajewsky, N. and Birchmeier, C. (2015a). Insm1 cooperates with Neurod1 and Foxa2 to maintain mature pancreatic beta-cell function. EMBO J 34, 1417–1433.

Jia, S., Wildner, H. and Birchmeier, C. (2015b). Insm1 controls the differentiation of pulmonary neuroendocrine cells by repressing Hes1. Dev Biol 408, 90–98.

Kristan, W. B., Jr. (2016). Early evolution of neurons. Curr Biol 26, R949–R954.

Kriventseva, E. V., Kuznetsov, D., Tegenfeldt, F., Manni, M., Dias, R., Simao, F. A. and Zdobnov, E. M. (2019). OrthoDB v10: sampling the diversity of animal, plant, fungal, protist, bacterial and viral genomes for evolutionary and functional annotations of orthologs. Nucleic acids research 47, D807–D811.

Layden, M. J., Boekhout, M. and Martindale, M. Q. (2012). Nematostella vectensis achaete-scute homolog NvashA regulates embryonic ectodermal neurogenesis and represents an ancient component of the metazoan neural specification pathway. Development 139, 1013–1022.

Layden, M. J. and Martindale, M. Q. (2014). Non-canonical Notch signaling represents an ancestral mechanism to regulate neural differentiation. Evodevo 5, 30.

Layden, M. J., Rentzsch, F. and Rottinger, E. (2016). The rise of the starlet sea anemone Nematostella vectensis as a model system to investigate development and regeneration. WIREs Dev Biol, doi: 10.1002/wdev.1222.

Lee, C. S., Perreault, N., Brestelli, J. E. and Kaestner, K. H. (2002). Neurogenin 3 is essential for the proper specification of gastric enteroendocrine cells and the maintenance of gastric epithelial cell identity. Genes & Development 16, 1488–1497.

Letunic, I., Khedkar, S. and Bork, P. (2021). SMART: recent updates, new developments and status in 2020. Nucleic acids research 49, D458–D460.

Liebeskind, B. J., Hofmann, H. A., Hillis, D. M. and Zakon, H. H. (2017). Evolution of Animal Neural Systems. Annual Review of Ecology, Evolution, and Systematics, Vol 48 48, 377–398.

Lorenzen, S. M., Duggan, A., Osipovich, A. B., Magnuson, M. A. and Garcia-Anoveros, J. (2015). Insm1 promotes neurogenic proliferation in delaminated otic progenitors. Mech Dev 138 Pt 3, 233–245.

Lutolf, S., Radtke, F., Aguet, M., Suter, U. and Taylor, V. (2002). Notch1 is required for neuronal and glial differentiation in the cerebellum. Development 129, 373–385.

Marlow, H. Q., Srivastava, M., Matus, D. Q., Rokhsar, D. and Martindale, M. Q. (2009). Anatomy and development of the nervous system of Nematostella vectensis, an anthozoan cnidarian. Dev Neurobiol 69, 235–254.

Mellitzer, G., Bonne, S., Luco, R. F., Van De Casteele, M., Lenne-Samuel, N., Collombat, P., Mansouri, A., Lee, J., Lan, M., Pipeleers, D., et al. (2006). IA1 is NGN3-dependent and essential for differentiation of the endocrine pancreas. EMBO J 25, 1344–1352.

Monaghan, C. E., Nechiporuk, T., Jeng, S., McWeeney, S. K., Wang, J., Rosenfeld, M. G. and Mandel, G. (2017). REST corepressors RCOR1 and RCOR2 and the repressor INSM1 regulate the proliferation-differentiation balance in the developing brain. Proc Natl Acad Sci U S A 114, E406–E415.

Moroz, L. L. (2009). On the independent origins of complex brains and neurons. Brain Behav Evol 74, 177–190.

Nakanishi, N., Renfer, E., Technau, U. and Rentzsch, F. (2012). Nervous systems of the sea anemone Nematostella vectensis are generated by ectoderm and endoderm and shaped by distinct mechanisms. Development 139, 347–357.

Nuchter, T., Benoit, M., Engel, U., Ozbek, S. and Holstein, T. W. (2006). Nanosecond-scale kinetics of nematocyst discharge. Curr Biol 16, R316–318.

Oliver, D., Brinkmann, M., Sieger, T. and Thurm, U. (2008). Hydrozoan nematocytes send and receive synaptic signals induced by mechano-chemical stimuli. The Journal of experimental biology 211, 2876–2888.

Pajcini, K. V., Speck, N. A. and Pear, W. S. (2011). Notch signaling in mammalian hematopoietic stem cells. Leukemia 25, 1525–1532.

Park, E., Hwang, D. S., Lee, J. S., Song, J. I., Seo, T. K. and Won, Y. J. (2012). Estimation of divergence times in cnidarian evolution based on mitochondrial protein-coding genes and the fossil record. Molecular Phylogenetics and Evolution 62, 329–345.

Renfer, E., Amon-Hassenzahl, A., Steinmetz, P. R. and Technau, U. (2009). A muscle-specific transgenic reporter line of the sea anemone, Nematostella vectensis. Proc Natl Acad Sci U S A 107, 104–108.

Renfer, E. and Technau, U. (2017). Meganuclease-assisted generation of stable transgenics in the sea anemone Nematostella vectensis. Nat Protoc 12, 1844–1854.

Rentzsch, F., Layden, M. and Manuel, M. (2017). The cellular and molecular basis of cnidarian neurogenesis. Wiley interdisciplinary reviews. Developmental Biology 6.

Richards, G. S. and Rentzsch, F. (2014). Transgenic analysis of a SoxB gene reveals neural progenitor cells in the cnidarian Nematostella vectensis. Development 141, 4681–4689.

Richards, G. S. and Rentzsch, F. (2015). Regulation of Nematostella neural progenitors by SoxB, Notch and bHLH genes. Development 142, 3332–3342.

Rosenbaum, J. N., Duggan, A. and Garcia-Anoveros, J. (2011). Insm1 promotes the transition of olfactory progenitors from apical and proliferative to basal, terminally dividing and neuronogenic. Neural Dev 6, 6.

Sebe-Pedros, A., Saudemont, B., Chomsky, E., Plessier, F., Mailhe, M. P., Renno, J., Loe-Mie, Y., Lifshitz, A., Mukamel, Z., Schmutz, S., et al. (2018). Cnidarian Cell Type Diversity and Regulation Revealed by Whole-Organism Single-Cell RNA-Seq. Cell 173, 1520–1534 e1520.

Siebert, S., Farrell, J. A., Cazet, J. F., Abeykoon, Y., Primack, A. S., Schnitzler, C. E. and Juliano, C. E. (2019). Stem cell differentiation trajectories in Hydra resolved at single-cell resolution. Science 365.

Steinmetz, P. R. H., Aman, A., Kraus, J. E. M. and Technau, U. (2017). Gut-like ectodermal tissue in a sea anemone challenges germ layer homology. Nat Ecol Evol 1, 1535–1542.

Stivers, C., Brody, T., Kuzin, A. and Odenwald, W. F. (2000). Nerfin-1 and -2, novel Drosophila Zn-finger transcription factor genes expressed in the developing nervous system. Mech Dev 97, 205–210.

Sunagar, K., Columbus-Shenkar, Y. Y., Fridrich, A., Gutkovich, N., Aharoni, R. and Moran, Y. (2018). Cell type-specific expression profiling unravels the development and evolution of stinging cells in sea anemone. BMC Biol 16, 108.

Tavano, S., Taverna, E., Kalebic, N., Haffner, C., Namba, T., Dahl, A., Wilsch-Brauninger, M., Paridaen, J. and Huttner, W. B. (2018). Insm1 Induces Neural Progenitor Delamination in Developing Neocortex via Downregulation of the Adherens Junction Belt-Specific Protein Plekha7. Neuron 97, 1299–1314 e1298.

Technau, U. and Steele, R. E. (2011). Evolutionary crossroads in developmental biology: Cnidaria. Development 138, 1447–1458.

Telford, M. J., Budd, G. E. and Philippe, H. (2015). Phylogenomic Insights into Animal Evolution. Curr Biol 25, R876–887.

Varoqueaux, F. and Fasshauer, D. (2017). Getting Nervous: An Evolutionary Overhaul for Communication. Annu Rev Genet 51, 455–476.

Watanabe, H., Fujisawa, T. and Holstein, T. W. (2009a). Cnidarians and the evolutionary origin of the nervous system. Dev Growth Differ 51, 167–183.

Watanabe, H., Hoang, V. T., Mattner, R. and Holstein, T. W. (2009b). Immortality and the base of multicellular life: Lessons from cnidarian stem cells. Semin Cell Dev Biol 20, 1114–1125.

Watanabe, H., Kuhn, A., Fushiki, M., Agata, K., Ozbek, S., Fujisawa, T. and Holstein, T. W. (2014). Sequential actions of beta-catenin and Bmp pattern the oral nerve net in Nematostella vectensis. Nature Communications 5, 5536.

Welcker, J. E., Hernandez-Miranda, L. R., Paul, F. E., Jia, S., Ivanov, A., Selbach, M. and Birchmeier, C. (2013). Insm1 controls development of pituitary endocrine cells and requires a SNAG domain for function and for recruitment of histone-modifying factors. Development 140, 4947–4958.

Westfall, I. A. (1996). Ultrastructure of synapses in the first-evolved nervous systems. J Neurocytol 25, 735–746.

Wildner, H., Gierl, M. S., Strehle, M., Pla, P. and Birchmeier, C. (2008). Insm1 (IA-1) is a crucial component of the transcriptional network that controls differentiation of the sympatho-adrenal lineage. Development 135, 473–481.

Wiwatpanit, T., Lorenzen, S. M., Cantu, J. A., Foo, C. Z., Hogan, A. K., Marquez, F., Clancy, J. C., Schipma, M. J., Cheatham, M. A., Duggan, A., et al. (2018). Trans-differentiation of outer hair cells into inner hair cells in the absence of INSM1. Nature 563, 691–695.

