## Supplementary Material for "*Insm1*-expressing neurons and secretory cells develop from a common pool of progenitors in the sea anemone *Nematostella vectensis*"

**SUPPLEMENTARY INFORMATION**

- **Supplementary figures and legends (Figures S1 – S7)**
- **Supplementary table S1**

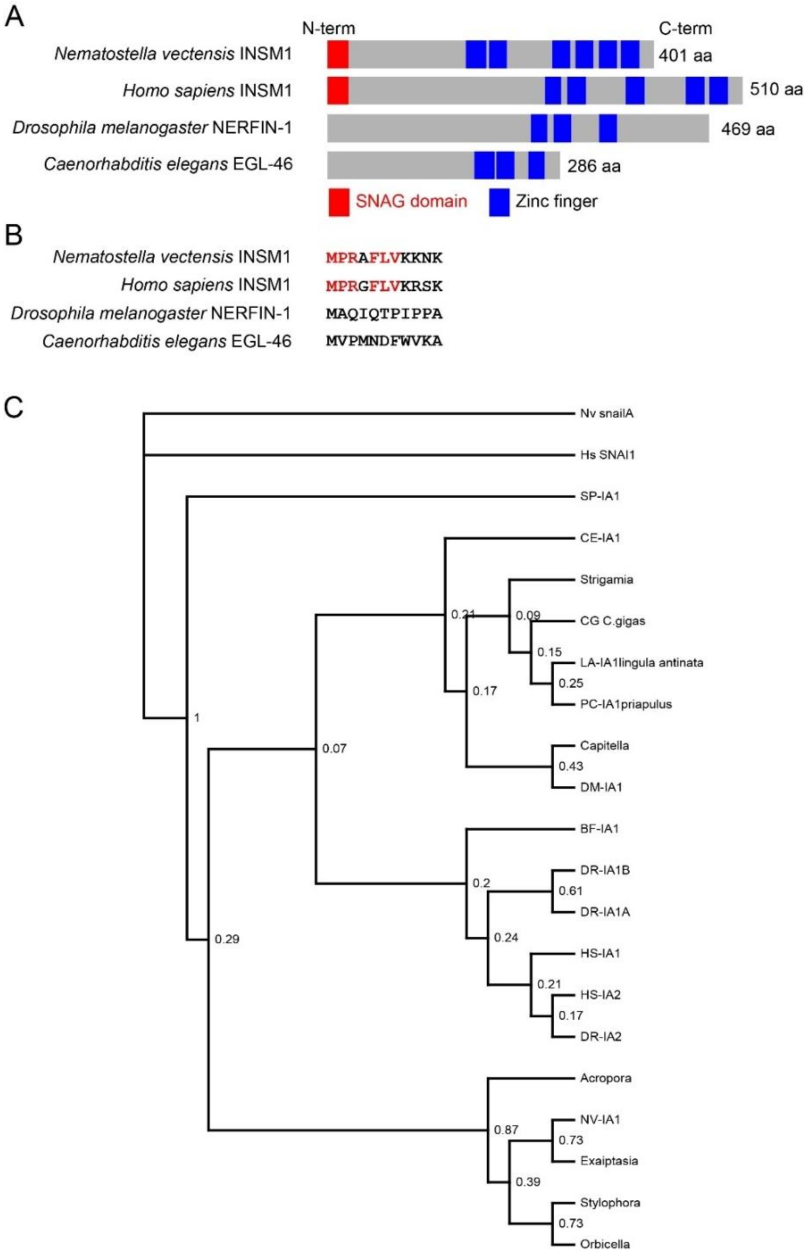


**Figure S1: Domain architecture and phylogenetic analysis of *Insm1* genes.**

(A) Schematic depiction of INSM1 proteins from different species. The domain architecture of NvINSM1 is highly similar to that of human INSM1 and in contrast to the *Drosophila* and *C. elegans* orthologs it contains a SNAG domain at its N-terminus. The amino acids of the SNAG domain are indicated in red in (B). (C) Phylogenetic analysis by Maximum likelihood (PhyML3.1) and based on amino acid alignments of Zinc fingers 1-3 (see Material and methods for details) and with *Nematostella* and human Snail proteins as outgroup. *Nematostella vectensis Insm1* protein is shown in bold and is placed in a clade with *Insm1* proteins from different cnidarians. Posterior probabilities are indicated at each node.

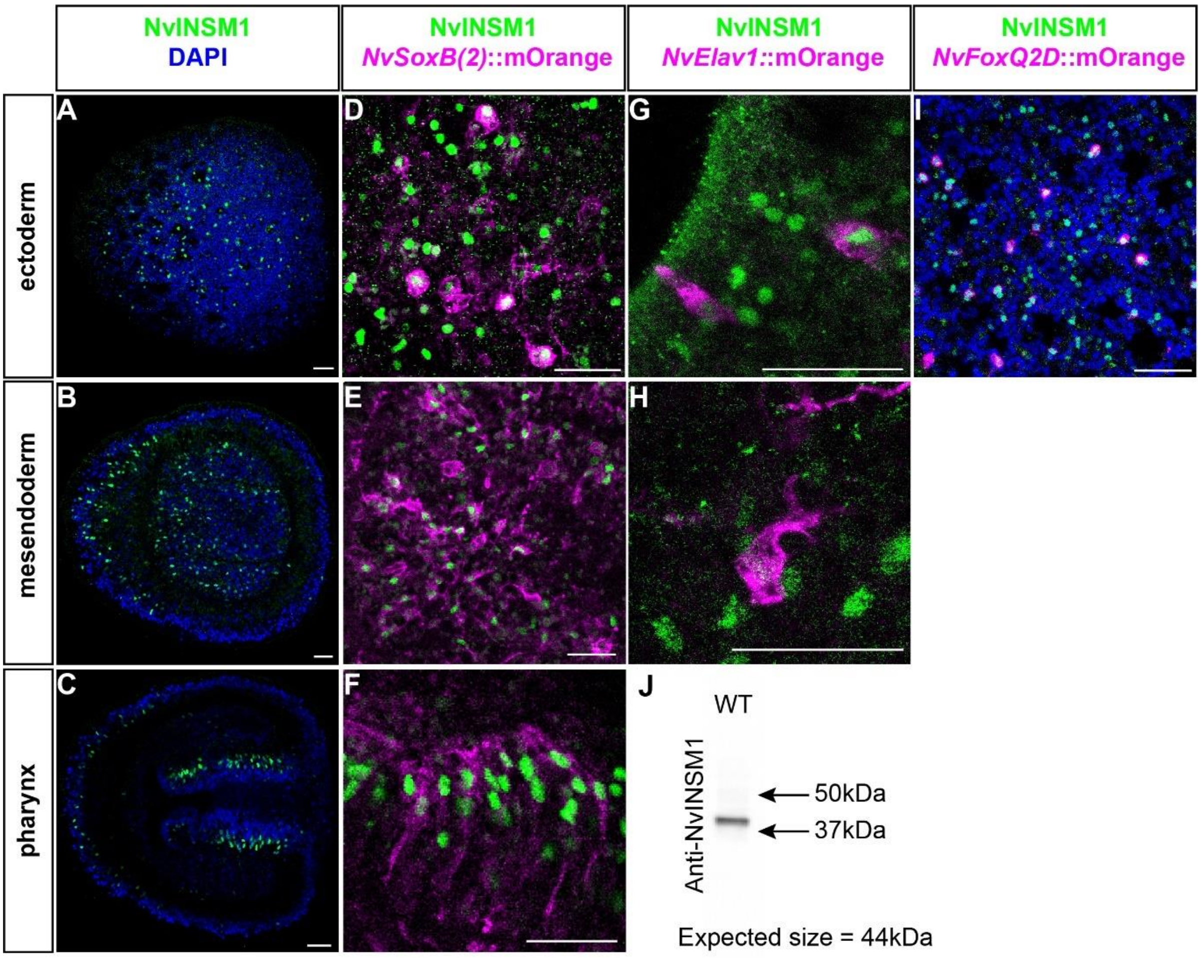


**Figure S2: NvINSM1 is present in various neuronal cell lineages**

Immunostaining of NvINSM1(green) and DAPI (blue) and various transgenic lines at planula stage (A-I) with staining indicated on the top. (A-C) lateral planula views (with the aboral pole to the left) at different stack level showing the presence of NvINSM1 in the ectoderm, mesendoderm and developing pharynx. (D-E) NvINSM1 staining within *NvSoxB(2)*::mOrange$^+$ cells (D-F), *NvElav1*::mOrange$^+$ cells (G, H) and *NvFoxQ2d*::mOrange$^+$ cells (I). Scale bars represent 20 µm. (J) Western blot on protein from wild-type (WT) animals. The blot was probed for NvINSM1. Arrows on the right indicate the positions of the ladder. The calculated mass of NvINSM1 44 kDa.

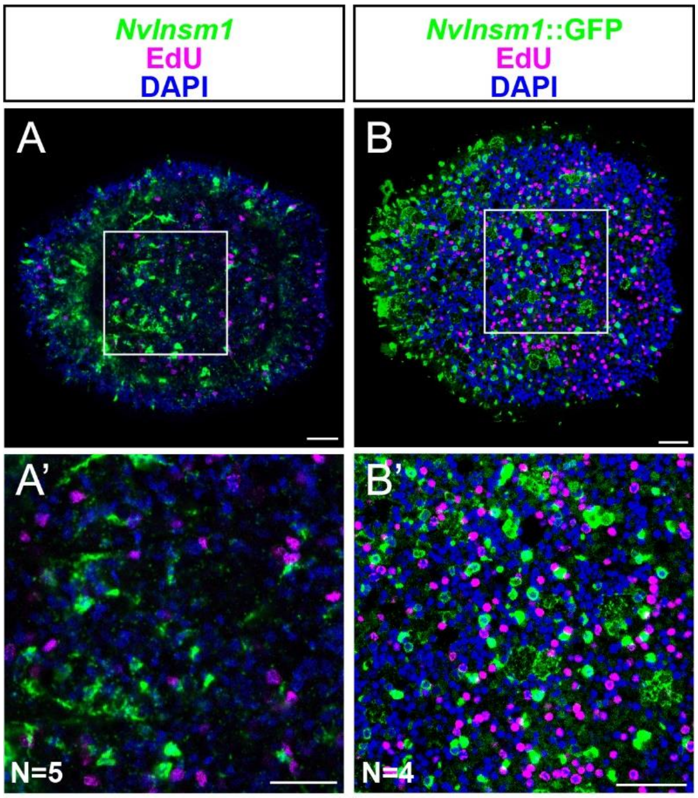


| *NvInsm1*$^+$ | *NvInsm1*$^+$ +EdU$^+$ | *NvInsm1*::GFP | *NvInsm1*::GFP +EdU$^+$ |
|---|---|---|---|
| 377 | 0 | 620 | 0 |

**Figure S3: *NvInsm1* expressing cells are post-mitotic.**

(A, A') Fluorescence in situ hybridization for *NvInsm1* (green) after 30 min pulse labeling with EdU (magenta). Lateral view with aboral pole to the left, (A') is a higher magnification of the boxed area in (A). (B, B') anti-GFP immunohistochemistry (green) in *NvInsm1*::GFP planula after a 30min EdU pulse (magenta). Lateral view with aboral pole to the left. DAPI is shown in blue. No *NvInsm1*/EdU double positive cells were observed. Scale bars represent 20µm.

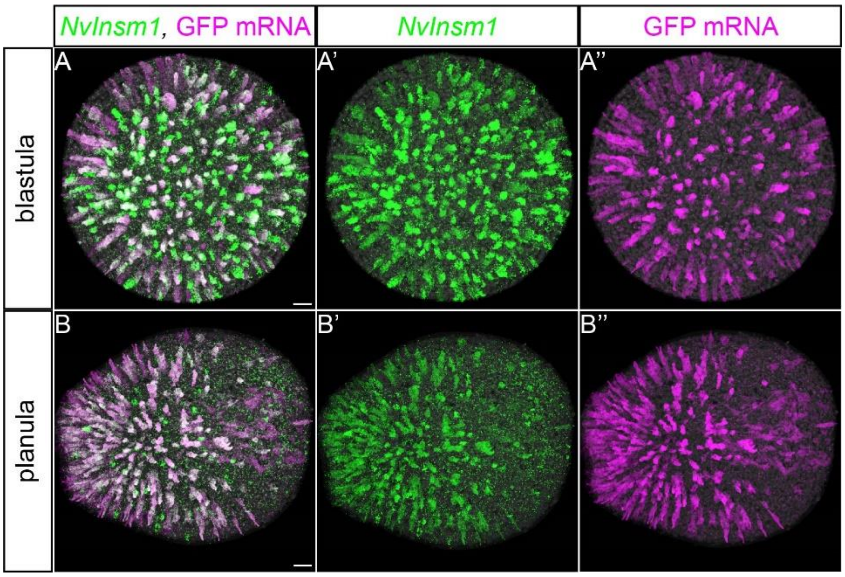


**Figure S4: Co-localization of *NvInsm1* and *GFP* mRNA in transgenic embryos.**

Double fluorescence *in situ* hybridizations, using probes for *NvInsm1* (green), *GFP* (magenta) and DAPI (grey), demonstrates that the reporter gene expression mimics endogenous *NvInsm1* expression at blastula stage (A) and at planula stage (B) in embryos from the *NvInsm1*::GFP transgenic line. Only few *NvInsm1*-expressing cells do not express *GFP*. Lateral views with the aboral pole to the left. Scale represents 20 µm

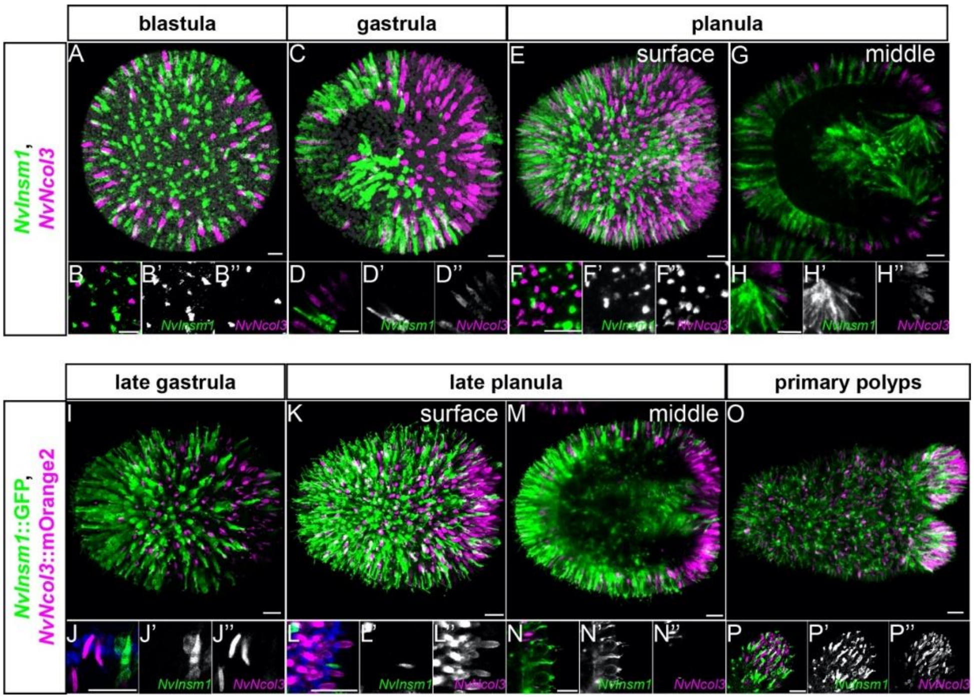


**Figure S5: *NvInsm1*-expressing cells do not give rise to cnidocytes.**

Confocal images of double fluorescence *in situ* hybridization (A-H) and double transgenics (I-P), developmental stages are indicated at the top and probes/transgenes on the left. (A-H) Double fluorescence *in situ* hybridization for *NvInsm1* (green) and the cnidocyte differentiating marker *NvNcol3* (magenta) shows no co-expression from blastula stage to planula stage. (B, D, F) represents ectodermal zoom regions at blastula, gastrula and planula respectively. (H) is a higher magnification of the pharynx. (I-P) Double transgenic animals with *NvInsm1*::GFP (green) and *NvNcol3*::mOrange2 (magenta) show that the two transgenes are not co-expressed. (J, L, N) are higher magnifications of the ectoderm at the different stages. (P) shows that even in the forming tentacles the two transgenes are not co-expressed, which suggests that *NvInsm1* is not expressed in differentiating cnidocytes. (A, C, E, I, K and O) are Imaris snapshots from the 3D reconstruction to show the overview. Some white spots on the overview pictures are due to the 3D reconstruction and do not reflect co-expression of the different labeling. All other pictures are stacks of two to three confocal sections to ensure that there is no co-expression. All pictures are lateral view with aboral pole to the left. Scale bars correspond to 20 µm.

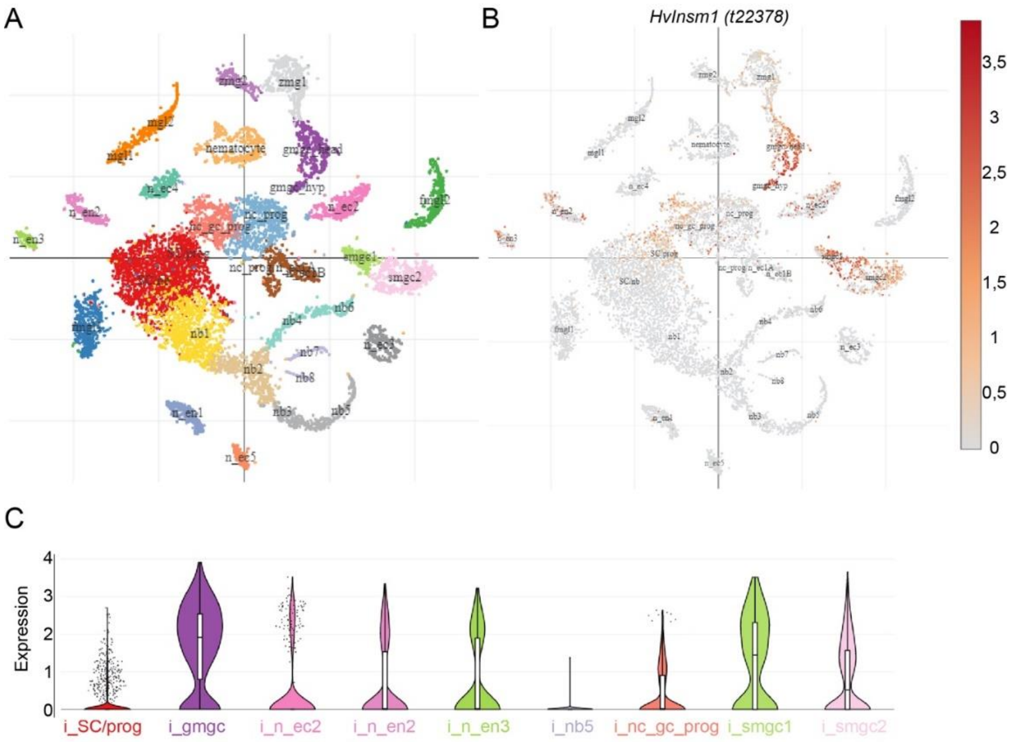


**i_SC/prog:** Interstitial, stem cell/progenitors

**i_gmgc:** Interstitial, granular mucous gland cells

**i_n_ec2:** Interstitial, ectodermal neurons 2

**i_n_en2:** Interstitial, endodermal neurons 2

**i_n_en3:** Interstitial, endodermal neurons 3

**i_nb5:** Interstial, nematoblast 5

**i_nc_gc_prog:** Intertistial, neuronal cells, gland celsl, progenitors

**i_smgc1:** Interstitial, spumous mucous gland cells 1

**i_smgc2:** Interstitial, spumous mucous gland cells 2

**Figure S6: Published single cell RNA sequencing data suggest that *HvInsm1* is expressed in neural and gland/secretory cells in *Hydra*.**

(A) tSNE plot of SNN (Shared Nearest Neighbour) clustered single cell data for interstitial cells mapped to the LRv2 transcriptome reference (Siebert et al.2019). (B) *HvInsm1* expressed in progenitor cells, ectodermal and endodermal neuronal cells and in various gland cells. Grey present low expression whereas red represents high expression (C) Violin plots showing *HvInsm1* expression in the different cell populations. The nematoblast 5 population (I_nb5) is here used as negative control. This analysis was done via the Broad Single-Cell Portal available at: https://portals.broadinstitute.org/single_cell/study/SCP260/stem-cell-differentiation-trajectories-in-hydra-resolved-at-single-cell-resolution

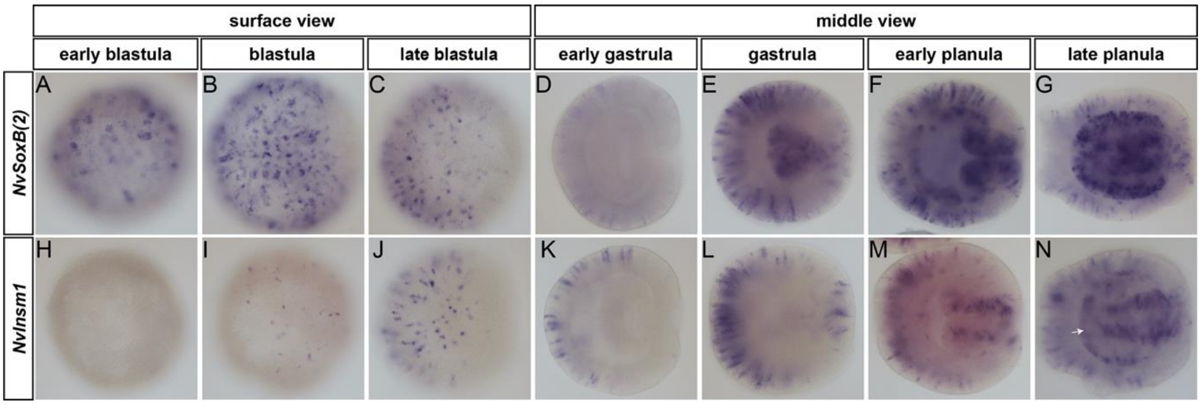


**Figure S7: Expression of *NvSoxB(2)* precedes that of *NvInsm1***

(A-G) represents published *NvSoxB(2)* expression pictures (Figure S1 in (Tourniere et al., 2020). The comparison with the onset of *NvInsm1* expression was performed during the same experiment, in parallel. *NvSoxB(2)* and *NvInsm1* are both expressed in scattered single cells all over the embryos, but the onset of *NvSoxB(2)* expression starts at early blastula stage (at 10hpf) before *NvInsm1* (12hpf). At gastrula stage, *NvSoxB(2)* is expressed in the forming pharynx and it's only at early planula stage that *NvInsm1* is also expressed there (E, M). At early planula stage, *NvSoxB(2)* starts being expressed in the mesendoderm, *NvInsm1* is expressed in this region at late planula stage (F, N) shown by the white arrow in figure N. All the embryos used for this experiment came from the same batch.

### Table S1

| Species | Gene | Accession number | Protein sequence |
|---|---|---|---|
| ***Homo sapiens*** | *Insm1* | NP_002187.1 | MPRGFLVKRSKKSTPVSYRVRGGEDGDRALLLSPSCGGARAEPPAPS<br>PVPGPLPPPPPAERAHAALAAALACAPGPQPPPQGPRAAHFGNPEAA<br>HPAPLYSPTRPVSREHEKHKYFERSFNLGSPVSAESFPTPAALLGGGG<br>GGGASGAGGGGTCGGDPLLFAPAELKMGTAFSAGAEAARGPGPGPP<br>LPPAAALRPPGKRPPPPTAAEPPAKAVKAPGAKKPKAIRKLHFEDEV<br>TTSPVLGLKIKEGPVEAPRGRAGGAARPLGEFICQLCKEEYADPFALA<br>QHKCSRIVRVEYRCPECAKVFSCPANLASHRRWHKPRPAPAAARAP<br>EPEAAARAEAREAPGGGSDRDTPSPGGVSESGSEDGLYECHHCAKKF<br>RRQAYLRKHLLAHHQALQAKGAPLAPPAEDLLALYPGPDEKAPQEA<br>AGDGEGAGVLGLSASAECHLCPVCGESFASKGAQERHLRLLHAAQV<br>FPCKYCPATFYSSPGLTRHINKCHPSENRQVILLQVPVRPAC |
| ***Homo sapiens*** | *Insm2* | NP_115983.3 | MPRGFLVKRTKRTGGLYRVRLAERVFPLLGPQGAPPFLEEAPSASLP<br>GAERATPPTREEPGKGLTAEAAREQSGSPCRAAGVSPGTGGREGAE<br>WRAGGREGPGPSPSPSPSPAKPAGAELRRAFLERCLSSPVSAESFPGG<br>AAAVAAFSCSVAPAAAPTPGEQFLLPLRAPFPEPALQPDPAPLSAALQ<br>SLKRAAGGERRGKAPTDCASGPAAAGIKKPKAMRKLSFADEVTTSP<br>VLGLKIKEEEPGAPSRGLGGSRTPLGEFICQLCKEQYADPFALAQHRC<br>SRIVRVEYRCPECDKVFSCPANLASHRRWHKPRPAAANAATVSSAD<br>GKPPSSSSSSSRDSGAIASFLAEGKENSRIERTADQHPQARDSSGADQ<br>HPDSAPRQGLQVLTHPEPPLPQGPYTEGVLGRRVPVPGSTSGGRGSEI<br>FVCPYCHKKFRRQAYLRKHLSTHEAGSARALAPGFGSERGAPLAFA<br>CPLCGAHFPTADIREKHRLWHAVREELLLPALAGAPPETSGPSGPSD<br>GSAQQIFSCKHCPSTFFSSPGLTRHINKCHPSESRQVLLLQMPLRPGC |
| ***Drosophila melanogaster*** | *Nerfin-1* | NP_524783.1 | MAQIQTPIPIPAPAEPIEKLSHPHSIGIQLQHKGMPHGLGLLTTVASLPP<br>KYRSRYLNLKTRCEIPLDLTLKLPSPLPSSDVPMAATFAEVYSENTSA<br>SSSEVQGEVTKVEELPVKPPTPPQSPAGKRKLSRESYDDEPAKMAKK<br>EEEVPVETVESTPAKPVKVAESAPKTNKASTSSGKPSRNKATRKLKF<br>DEETSSPVSGTVIRPLEDITDGSMQYSNGDIDPKYNIVEITEETKAELA<br>AIKNVIGDYVCRLCKIKFEDAFGLARHRCACIVLLEYRCPECGKQFN<br>CPANLASHRRWHKPRKEASKKENRNTTNQPEKQQQVEKKSEEELAF<br>DCQECGKKFKRAAYLRKHQLTHQKKEKPAEKQLEMKPTTTTITGSF<br>HFNQAVSTSSSASSSHSDEGVYYVVGSNARENYDYDNDYLSDSSSSAS<br>SAGYGRLQIVEHGLTEEESIAAAALTNLRNCASVIQHTTMAH |
| ***Caenorhabditis elegans*** | *Egl-46* | NP_504694.1 | MVPMNDFWVKAILSSTNPSPVPSTTSTVSNDENLDKTLDFDCSTQTV<br>FPTLPMFWNPTLVQQMLALYQIQQQQIQFSAKLAPQPLFQEPTIQKEF<br>LPFPHQSRKRPLPIDPKKTKLRKLNEDTVTSSPVSGMFIKEEADVKSV<br>EELQKEADLLDETAAYVEVTEESRQKIDEIPNVIGDCICRLCKVKYED<br>VFKLAQHKCPRIAHEEYKCPDCDKVFSCPANLASHRRWHKPRNELG<br>GSPPAQSSTIVSCSTCFNSFPTKKMLKLHSSTCQRSPLQDLLSRVIPTM |
| ***Danio rerio*** | *Insm1A* | NP_991207.1 | MPRGFLVKRNKKATPVSYRVRSEEDEQGAFVAQDVPCARRPVSPVQ<br>FGNPETVYRAMYSPTRPVSREHERACLERRFNLGSPISAESFPAAPNC<br>SDQAPVDLKIGTSNSNRTGTTVTTKRPASDTERKGKPASKKAKAMR<br>KLQFEDEMTTSPVLGLKIKEGPVEQKPRSQCASGDKPPGEFVCQLCR<br>EAYADPFSLAQHKCSRIVRIEYRCPECDKLFSCPANLASHRRWHKPK<br>QSAESNKTPAPEKEETSSDRDTPSPGLSESGSEDGLYDCQHCGKKFK<br>RQAYLKKHVTAHHDAPEKPQSHAPLNLSASECHLCPVCGENFPSRM<br>SQERHIRLQHSAQVYPCKYCPAMFYSSPGLTRHINKCHPSENRQVILL<br>QMPVRPAC |
| ***Danio rerio*** | *Insm1B* | NP_955952.1 | MPKGFLVKRNKKAALISYRIRTDEDGGPTPECPIAQIALSSPAPSASKP<br>DSILLAFPSAGAEAPVPVHKPVQFGNPEAVYQALYSPTRPVSKDHDR<br>KYFERSLNLGSPISAESFPTPASLTSLDHHLLFAPVDLKIGTSNSNRSG<br>TASGAHAPAIRTGAKRPSADAAERKVSSKSAKKPKAIRKLNFEDEVT<br>TSPVLGLKIKEGPVDLKPRPSSGGTNKPLGEFICQLCKEEYSDPFSLAQ<br>HKCSRIVRVEYRCPECEKVFSCPANLASHRRWHKPRVQSAPKQALQP<br>AKPFPEELRAEFPSDRDTPSPGLSESGSEDGLYDCQHCGKRFKRQAY<br>LRKHILGHQALQNQILGEAFRSAESPDAMPSEDRQSPAPLNLSPADCL<br>TCPACGEKLPNRASLERHLRPLHDDAQAFPCKFCPATFYSSPGLTRHI<br>NKCHPTENRQVILLQMPVRNAC |

| | | | |
|---|---|---|---|
| ***Danio rerio*** | *Insm 2* | XM_001332 478.8 | MPRGFLVKRSKRGSSASYKVRAEETEPRKEDLNTQTPNASVLEPIRES WTSEISSKHEETRLLEDAGGSADYAQNYFIHPEQSASSPRNTCDSYSPI KPISTELLDRCLSSPAMAESFPLVTPVSSIERLLMNHSDMKFGTPVPSS VPTYPALHQCVKRAFMDSERKSKPPKKPKVIRKLNFEDEVTTSPVLG LKIKKESPETKLPPQGGRKKPLGEFICQLCKEEYPDPFSLAQHKCSRIV RVEYRCPECDKVFSCPANLASHRRWHKPRTLSTADTKKSQLEARNL EKTSSVEGKENASKMRVNNQHQLSPDSSQLHRSAPDSSMMHRDPPL DQRFSTPEKCFEMHIGSAEASRLFDHCPEEPDRSPATPYLPSPGPEEVY DCHYCSKKFRRQAYLRKHLAAHETIKASSYGQIESGQITFPCHLCGA HFPSAEIRDKHRLWHAVREDLLLRPDLSPAAGDQQIFSCKHCPSTFFS SPGLTRHINKTHPSESRQVMLLQMAVRPGC |
| ***Strongylocent rotus purpuratus*** | *Insm 1* | XM_003728 770.2 | MPRNFLVKRTKRTGSVTYRQRTSDEEYDDLGFVDGDLTDAYIRPFNS PDSGYSQSPVAPISHKDPSLAFQFDRDATVRSPNVLQASGPALTSSHIP LTPYFTTGYKASMPSPVTPNIKLTLSAPPVLGTKRPNSDSDHKPHRAK KPKAARKLTFEEDNRSPVHGTIIREEVDITSKTNSLGKTSDQNGTKLG GDFLCKLCKESYSDPLSLASHKCSCIVHVEYRCPECDKVFNCPANLA SHRRWHKPKPVNNTSNSQRSTPTNSPRILPAGGAKEALKHYPTTVLN IPDTTRKGDSGSEGSMSRGSTPSPLHHLFPTSMDTPRSTPSPSSTYNNN NNDTKTSHGGDMLYHCDQCGRKFRRPANLRRHMQQHGEEETFPCQ YCGRVFNSLTSRAKHVLTHAVSNNNNSNTATKPSNRDCEDNYDQRV ALDRLEKLQQHGLPGEMYACKYCASIFPTSPGLTRHINKMHPSENRQ VILLQMPALRT |
| ***Branchiostom a floridae*** | *Insm 1* | XM_002610 125.1 | MPRGFLVKRGKKFVPVSYRQRDEDGVATQKMEPPASPVQRHPCQPA YPDSPHFTYSPIRPVTREADSTLRESFGYHKSSSPLSAHAFPSPLMFDQ YSFSPASGGPREKYSDTPSPNSGSAKRPHPDGERKPKSPIKKSKAVRK INFDEDTTSPVLGLRITQAPAEQEKQKDKDKGDSNNNSSKTTNSTNK KPGGGAFVCQLCKEEYTDPFTLAQHKCSRIVRVEYRCPECDKVFNCP ANLASHRRWHKPRPTESPNSRFSAASSPSRTHSDSGSHRSTPSPHTDA SHHRDSVHADTPWFECDRCGKKFRRQAYLRKHILQHEEGKEEPSYP CHLCGKDHCCPECGAGFPNKAALDRHVRMHSSDIFSCKYCDATFYS SPGLTRHINKCHPSENRQVILLQPVPARHTPTAVC |
| ***Priapulus caudatus*** | *Insm 1* | XM_014807 664.1 | MPRGFLVKRRARPHVCVSYRAASVSDDDRSDDRSDSDDHTDVGSQS AWSPEPARNGNWNVGRPRDENAHQAAWSPEPPAGGRAWMPTLCRP DKWSPELWRAARSSPIRVPLYRGGGGGGDSVSDRDSPLSLDSRSPGS TASSSPLPLSIHPYFTAGTSPLLYINGFDKLSVHSPGNMPATPKRATPT RPDNSSSPSSMSSPSKRRLVGDHGKPRTPKKSRAARRLCLDDETASP VSGTIIRAARDSDDAAADNNAGDDIVVEKCADIDAALNLVAMTPEA CADLAKIENRIGAYVCQLCKHRYEDAFQLAQHRCSRIVYIEYRCPEC DKVFNCPANLASHRRWHKPKPEATSPPREEKSDGGDAPREEQRRLEE TTCNSEGEYRCGTCGKTFKRQAYLRKHMQAHHVGGAPATPPRRPPT FDGGADRVRHVLALHGGAAGAALRQHRCIVCGCGFGTRAELDKHA RTHVSDTYPCKYCDSTFFTSPGLTRHINKCHPTENRQVILLQLPTI |
| ***Lingula anatina*** | *Insm 1* | XM_013544 550.1 | MPRGFLVKRHYQCGSAGGLQAHQLETSRLRRNSEEDRSDSGSEHDY VSHDTAQSPDSGFSASPLALTTRDRSASPGTRETPPEPRGSQLRPLSSL QPYYYVPSNLSYYHNTASTYSPFYFSSFDRLSVTSPTGKNIVTDSVVPQ CPNKKRSNDSISKPKPNRKSKAARRLNFSEVKNSPVSGTFIRDSDDED TGGDIGTSRVVRGDIDGSLNFVEVTEEAKNELAKIENKIGDYICQLCK EWFEDAFRLAQHRCSRIVHIEYRCPECDKVFNCPANLASHRRWHKPR PNSSTKVSAPSKILPAPPAAHDHDTPLNLAVKDRVNPSLGTPATTAES EALYSCDKCDKKFRRRAYLVKHLQTHQVESSEPAPRRVEPERPYPCR YCGQVLPSEEYLAKHLLQHNDEVSVIPSDRPRPPPSHLPNGTPCVQPD FLYKCKHCTSVFYTSAALTRHINKRHPSDTKQVILLNLPTPEMNRQ |
| ***Capitella teleta*** | *Insm 1* | tr\|R7UCX3\| R7UCX3_C APTE OX=283909 GN=CAPTE DRAFT_219 688 | MPKGFLVKRHKPLNLSTFRRRGSEEDPSDSERSASGSEHEDQHTSSSA PKILCPLLPTDNAQYSCSPDSGYANSPGSILPIKDDNENLAPRRNLDK CDISPGSGASHSRTSSPSFSPLSVHTPASFPHGFQFSAFDRLRVCSPQH RIGPAGPLFPGLHLLPHAAFHSGNYFPPPLNGNFGLPPSSLHAPLLMP APHSPFKAASRILSPVNPSQGSRPSPLLSNAYSMSTSPGKKRTASESISS ADMLNANSKPMKSAKKTKAQRKIHFDEDNSSPVSGTIIRDAPNEQNG FKVCSGDIDSSLNLVEVTPEARAELEKIDNKIGDYICQLCRDKFEDAF QLAQHKCSRIVHVEYRCPECDKVFNCPANLASHRRWHKPRPNGSTA QSKNPAAPKRAPSSPLSKPTDVVNGNYGMFMHEGDARGGSPGGSSS GSSDQGTDAPFECQACHRRFKREAYLRKHEATHHAPMHSPQSITERY SCHYCGESLKTEVARTKHILQMHTPGLGGLHQISRPAMPPMLPRSPE QVYRPEQAMHCKFCPSVFFSSPGLHRHISKCHSSESRQVMSYFFLPFF FIRVALHSGIL |
| ***Crassostrea gigas*** | *Insm 1* | XM_011433 622.2 | MPRGFLVKRTSHGGAASYRERRTSEEDRLDLEADMTRFGSPDSGYC ASPINFSKDIENNNYEKPRSSPLSVSTQQLYFSSFEQLSVNSPNNENSA |

| | | | |
|---|---|---|---|
| | | | PKPVDQSSPNKRRNENKNKPMKKPKAARKINFDEDTTSPVSGCIIKHVSPEDEGTVIYHGDIDSTYNFVEATPEARAELERIENKIGDYICQLCKEFYEDAFQLAQHRCSRIVHVEYRCPECGKVFNCPANLASHRRWHKPRPSPASSRPSAPAKILPSNQNENNANTMEPKNDNSMFLVPTDLSVSNAEGQFKCETCGKKFRRQAYLRKHRQVHNDSGQIARNDNSTRPEILHISNMNELSNKESLRRDVVSPQNAFESLQTRVADMYVCKYCESSFSSSPGLTRHINKCHPSENRQVILLQMPSTLPRHM |
| ***Strigamia maritima*** | *Insm 1* | tr\|T1JGD0\|T1JGD0_STRMM | MPRGFLVKRLPSSKHSAPSSSYRVRRYSDEDRSDSSSDHDNIAPPTQLPLPTTGGAQQLVAPAAHASPDSGFGSPTLLLYRSPTQFNSRTTNHFYNEKNGENSPLALTTRATSLATSLTTSLATSAVPYLNPFDQFMVSLCTPPIKSSQTPAPPPPSPHKRRESAVALPKGSRAPKKSKAVRRLTFDEDKSSPVSGTIIRDFDDPDEPQNTSLMVVRKGDIDPSFNLVEATEEAKAEIAKIENRIGDYICQLCKKCYDDAFGLAQHRCSRIIHVEYRCPECDKVFNCPANLASHRRWHKPRPNSAVQPPTSRVTAINNNNILPPPPRLLPSLQSLQSLQSLQGDLSDSESNSSESQCSPNREGVDEVFECDLCNKKFRRQNQLRRHLLTHSADKTLSDDIGADVGGDVGGSFACKFCPNAFATSPGLTRHINKLHPSETRQVILLQVPQH |
| ***Hydra vulgaris*** | *Insm 1* | Sc4wPfr_57.2.g24939.t1 | MGIKKVIMSLFLFFIEDIVFSVSTKPGASLSLVLKQNNKKNQIVKFRHYKKSRLFQPYMVLNNKSKPFRKDKTDLESVVALSMKTVFVKKLPSDKTEEIFTKDNKYSPSKENVVEGTSGKSLLLQAGFKCQLCGEYHSNALSLAYHKCSSIKHIEHRCPECNKAFSCQANLASHRRWHKPKIASTKRHNKNSPDPFKKSDRRQAVVIGNYKSKERLVKLGVGQGTCIGSLLFKIYVYDLSSATELDTYFFANDTTIVAKADSIEEYQELCCKELEKISNCFVANGHTLHPEKTKIMIFGSKKKSRSKVARSQGHPMW |
| ***Orbicella faveolata*** | *Insm 1* | XP_020618957.1 | MPKAFLVKKNKRSYVEASASEKRTLEYDAVPAYRELDSVVLSTASAVSDIMHSSTANSDKAPVLTSLSSLLTSTGTANGSFPSSAFKPFKVEKEKENSPAKKLKPNPPPLVESRDKNNRSKDNIGGREALNERHTDHGGKTAVLEHKEFAISDKYDFPKPKGPLPQSQFICQLCQEVYSDPFSLAQHKCSGIQHTEYRCPECDKVFSCPANLASHRRWHRPRSPHEKKATPTQTPAQREPQLPNSPKTSAEDNGKNGAVTTAVESEARSSSPVGNSQGQFVCEVCNKTFRRKAYLRKHMNNHNDDRPYPCQYCGKVFRSLTNRAKHVLNHAVGPKTFICNVCGNGFANKGSLDRHARIHTGEIYSCKHCSSTFYSSPGLTRHINKCHPPENRQVIVLQVPVRQG |
| ***Stylophora pistillata*** | *Insm 1* | XP_022777808.1 | MPKAFLVKKNKRSHGGTSVSERRTLEYDALPSYREGKSVELSTSSAVSDIMHSSSLANSDKAPVLTSLSTLLTSTSNGNVTSSAFKPFKVEKEKENSPKKFKPNPPLAENRDKPTRGRESNSSTGREALNEKHSDHGGKTAVLEHKEFAIYQDKYDFPKPKGPLPQSQFICQLCQEVYSDPFSLAQHKCSGIQHTEYRCPECDKVFSCPANLASHRRWHRPRSPHEKKPAPSQAPASRSEQLPNSPKTSGEDNKNGVTTTTESEVRSTSPIGNSQGQFVCEVCNKTFRRKAYLRKHMNNHNDDRPYPCQYCGKVFRSLTNRAKHVLNHAVGPKTFICNVCGNGFANKGSLDRHARIHTGEIYSCKHCSSTFYSSPGLTRHINKCHPPENRQVIVLQVPVRQG |
| ***Acropora digitifera*** | *Insm 1* | XP_015758892.1 | MPKAFLVKKNKRSHAGNGSTSEKRTLEYNPLPEYREIESAVHSTTSAVSEIMHSSPTKTAKVPVLTSLTTLLSSAPNGKANSTSAFTPFKVDKEKENSPAKKFKPGPTNIETEDKARKARDSNRATGRETLHEKHTDHGGKTAVLEHKEFAICQDKFDFPKPKGPLPQSQFICQLCQEIYSDPFSLAQHKCSGIQHTEYRCPECDKVFSCPANLASHRRWHRPRSPHDKKPAAAQTPTPRDQPASTNTKPSVEGSKIGVAATDSDDCTRAASPVGNSQGQFVCEVCNKTFRRKAYLRKHMNNHNDDRPYPCQYCGKVFRSLTNRAKHVLNHAVGPKTFICNVCGNGFANKGSLDRHARIHTGEIYSCKHCSSTFYSSPGLTRHMNKCHPPENRPVIVLQVPVRQG |
| ***Exaiptasia pallida*** | *Insm 1* | XP_020901918.1 | MPRAFLVKKNRSPRTAAGEKRSFPEDDYSPNFNYIDNRIFSTSSAVAGLIHSPTESDKFPELSSLTSMFTSTNTTASCNSAFKPFKPEKGLSPKKHKPTQPVTEVKSEKTKKNSKENTTVNLINERHSSEHGGAKTAVLEHKEFAIFSENSDFPKPKGPLPQSQFICQLCDEEYSDPFSLAQHKCSGIQHTEHRCPECDKVFSCPANLASHRRWHRPRSPHEKKSTSTTPQKETKKFDSVSESEGGQKSSSPPLESERSTSPVSNSQGQFVCNLCNKTFRRKAYLRKHMNNHTDDRPYPCQYCGKVFRSLTNRAKHVLNHAVGPKTFTCNVCGNGFANKGGLDRHSRIHTGEIYSCKQCSSTFYSSPGLTRHINKCHPPENRQVIVLQVPVRQG |
| ***Clytia hemisphaerica*** | *Insm 1* | TCONS_00054478 | MPKSFLIKKWGRKFNFYSRSSEFNIQKDCTEETNFLTIPSPPPARPVLKEKNLSSPTQPSNVKEKPIDLKALFLTSIPVSKPFGLYYLEQNSRSNNGQQPVTKRHVPPEDNPTTIIPSPNKDHIFRVSPKVDGPLNQSSFNCQLCKLSFCDPLSLAQHKCSKIKHVEHRCPECDKVFNCPANLASHRRWHRPRSPTTNRPRKVGKAEKKLVEKKSQISLRTPEYQIKNPKIEAMRNIREDPKLQEVPVRGSPIVKPTVYRYKQTTQNLPNRFYDQMNIYRQRKLSNHFPTMISNNRMMVRKHYHRIGSESSDVESEEDFLEHKQNHELSTEEIEIDV |

| | | | |
|---|---|---|---|
| | | | ISNNSNSSSMTSPSSSLTSPKLPYQQGLFKRGSWNGPSHTTPPPYTFSY<br>M |
| ***Nematostella vectensis*** | *Insm 1* | XP_001627769.1 | MPRAFLVKKNKSVKSSGEKRSYPEDDYDPNFNYVDNRIFATSSAVTG<br>IIHSPTECDKFPVLSSLTTMFTSTTASTSCTSAFKPFKTEKENSPKRHKS<br>SVHSTEGKAEKPSKRESKMEFSPISRDLHNEKLTNEHGSVGGKTAVL<br>EHKEFAIFSDKNDFPKPKGPLPQSQFICQLCSEVYSDPFSLAQHKCSGI<br>QHTEHRCPECDKVFSCPANLASHRRWHRPRSPHDKKSSQPQKPVEK<br>KKVESESESDSESDTEKKPGSPAVESELSPVGSSQGQFVCTVCNKTFR<br>RKAYLRKHMNNHTDDRPYPCQYCGKVFRSLTNRAKHVLNHAVGPK<br>TFTCNVCGNGFANKGGLDRHSRIHTGEIYSCKQCSSTFYSSPGLTRHI<br>NKCHPPENRQIVVLQVPVRQG |
| ***Homo sapiens*** | *Snail 1* | NP_005976.2 | MPRSFLVRKPSDPNRKPNYSELQDSNPEFTFQQPYDQAHLLAAIPPPE<br>ILNPTASLPMLIWDSVLAPQAQPIAWASLRLQESPRVAELTSLSDEDS<br>GKGSQPPSPPSPAPSSFSSTSVSSLEAEAYAAFPGLGQVPKQLAQLSEA<br>KDLQARKAFNCKYCNKEYLSLGALKMHIRSHTLPCVCGTCGKAFSR<br>PWLLQGHVRTHTGEKPFSCPHCSRAFADRSNLRAHLQTHSDVKKYQ<br>CQACARTFSRMSLLHKHQESGCSGCPR |
| ***Nematostella vectensis*** | *Snail A* | AAT72903.1 | MPRSFLVKKTCDKKALLLRNGPVMTREEGGKSVESHRKGSSPERVE<br>NEDKADKRLQYWNGLGTSLVYPAEVTSVPGLHNKPTEFCIDVKYER<br>SNTKDDVDMGLSRELEYGTKTASLTAEESRPKHQCHQCNKGYSTPL<br>GLAKHQQFHCNTHHKKSFTCKHCDKIYVSLGALKMHIRTHTLPCKC<br>SICGKAFSRPWLLQGHIRTHTGEKPYQCTNCKRAFADRSNLRAHMQ<br>THAVVKKYSCSRCKKSFSRMSLLVEHEDSGCPSQG |

**Table S1: Amino acid sequences of *Insm1* genes in various metazoans.** Accession numbers and amino acid sequences for the species included in the phylogenetic analysis (**Figure S1**) (retrieved by BLAST searches at NCBI and via OrthoDB (https://www.orthodb.org/)) (Kriventseva et al., 2019). *Clytia* and *Hydra* sequences are listed, but were not included in the phylogenetic analysis, because they encode only two zinc fingers.
